# Spiroplasma impairs testes gene expression in Glossina fuscipes fuscipes

**DOI:** 10.1101/2025.09.23.678099

**Authors:** Riccardo Piccinno, Giulia Fiorenza, Francesco Lescai, Simone Carpanzano, Fabian Gstöttenmayer, Kiswend-sida M. Dera, Anna Cleta Croce, Chantel J. de Beer, Mariangela Santorsola, Giuliano Gasperi, Federico Forneris, Adly M. M. Abd-Alla, Serap Aksoy, Anna Rodolfa Malacrida

## Abstract

*Glossina fuscipes fuscipes* is a riverine tsetse fly species, the primary vector of human and animal trypanosomiasis in Sub-Saharan Africa. Controlling tsetse fly populations is crucial for mitigating the socio-economic impact of this disease, as effective treatments remain challenging. Indeed, the development of control strategies is hindered by the species’ unique reproductive biology: adenotrophic viviparity, in which the female retains and nourishes the developing larva in her uterus throughout the pregnancy. The discovery of *Spiroplasma* in some *G. f. fuscipes* populations has drawn attention as a potential tool to enhance tsetse fly control strategies. Although *Spiroplasma* does not exhibit in *G. f. fuscipes* the male-killing phenotype observed in *Drosophila melanogaster*, evidence suggests that it may confer refractoriness to *Trypanosoma* infection. This has led to further investigations into its broader effects on *G. f. fuscipes* biology, particularly its potential impact on *Glossina* reproductive fitness. In this study, we considered *Spiroplasma* effect on the male reproduction. For this, we performed a differential gene expression analysis on testes and male accessory glands (MAGs) between *Spiroplasma*-infected and uninfected males. A significant downregulation of genes was observed in testes while a minor effect has been detected on MAGs. Downregulation of testes genes associated with functions related to sperm motility, energy metabolism, and mitochondrial function has been observed. Additionally, differentially expressed genes involved in antimicrobial activity and circadian rhythm regulation were observed. These findings provide valuable insights into the potential fitness costs of *Spiroplasma* infection for the fly and its implications for the bacterium use as biological control strategies targeting *G. f. fuscipes*.

## Introduction

*Glossina fuscipes fuscipes* (*Gff*) is a riverine species of the *Glossina* genus, comprising flies known as tsetse flies. In this genus, both sexes are obligate hematophagous, feeding on vertebrate hosts. These flies represent the primary vectors of African trypanosomes, the causative agents of African trypanosomiasis that affects both humans and animals. This disease is better known as sleeping sickness in humans and nagana in animals. It is presenting severe socioeconomic consequences, including fatal outcomes in humans and a reduced livestock productivity (Büscher *et al*., 2017; Rotureau & Van Den Abbeele, 2013). Infection occurs during the blood feeding when an infected fly injects the protozoan parasite into the host’s bloodstream (Rotureau & Van Den Abbeele, 2013; Wamwiri & Changasi, 2016). *Gff* is reported to be the main vector of Human African Trypanosomiasis (HAT), especially in Uganda (Krafsur, 2009). As such, it is the key target for vector control programs in endemic areas (Hyseni *et al*., 2012; Ozioko *et al*., 2020).

Several strategies have been developed for *Gff* populations control. In addition to traditional methods, such as traps and insecticides (Mbewe *et al*., 2018; Ozioko *et al*., 2020), this species is a target for biological methods such as the Sterile Insect Technique (SIT). This approach involves the release of sterilized males which may compete for mating with wild fertile females. The outcome of these sterile matings is a reduction of population size (Dera *et al*., 2025; Mahamat *et al*., 2023; Vreysen *et al*., 2000). The employment of parasites or symbiotic bacteria able to interfere with fly reproduction or with *Trypanosome* infection is now under consideration (Alam *et al*., 2011; Symula *et al*., 2013; Wamiti *et al*., 2018). A constraint for the planning of the new control approach is the peculiar reproductive strategy of this fly. Unlike most insects, *Glossina* species are viviparous, meaning females don’t lay eggs but give birth to fully developed larvae (Benoit *et al*., 2015). This type of reproduction includes different adaptations, among them the use of a spermatophore for the ejaculate transfer from the male to the female. Indeed, during mating, males transfer spermatozoa along with seminal proteins which interact with female-driven proteins to form a protein structure called spermatophore inside the female uterus that encapsulates the ejaculate (Attardo *et al*., 2020; Fiorenza *et al*., 2025). Along with spermatozoa, the ejaculate contains non-sperm components such as seminal fluid, proteins and carbohydrates which are produced in both testes and MAGs. In *Glossina,* species ejaculate displays plasticity because it plays a vital role in post-copulatory sexual selection and as a consequence, drives rapid evolution exchanges and rapid isolation (Scolari *et al*., 2016; Savini *et al*., 2021). *Wolbachia*, a well-known bacterium capable of manipulating insect reproductive behaviour, by causing cytoplasmic incompatibility (CI), has already been employed in the biological control methods for sanitary and agricultural pests (Gong *et al*., 2023; Montenegro *et al*., 2024; Werren *et al*., 2008). However, a low titer of *Wolbachia* has been detected in geographic populations of *Gff*. This opens the question about the possible use of *Wolbachia* as a control method (Doudoumis *et al*., 2017; Symula *et al*., 2013).

In 2017, *Spiroplasma* was identified in *Glossina* species belonging to the Palpalis group, which includes *Gff* (Doudoumis *et al*., 2017; Schneider *et al*., 2019). The presence of this bacterium has different effects among different species: from beneficial to pathogenic (Arai *et al*., 2022; Ballinger & Perlman, 2019; Bolaños *et al*., 2015). A *Spiroplasma* belonging to the Citri clade, Poulsonii group, was identified in the hemolymph of a laboratory line of *Gff* (Fiorenza *et al*., 2025) and has been characterized as *s*Gff (Bruzzese *et al*., 2025). This strain, by contrast to *S. poulsonii* in *Drosophila melanogaster*, does not exhibit the male-killing pathogenic effect (Arai *et al*, 2022). A very important feature of *s*Gff is that it is able to render *Gff* refractory to trypanosome infections (Schneider *et al*., 2019). This characteristic makes it an interesting potential tool for implementing disease control. As a consequence, analysis on the impact of *s*Gff on the reproductive biology of *Gff* is necessary to determine its suitability as a control agent.

Preliminary analyses have shown that *s*Gff can be transmitted vertically from mother to larvae (Son *et al*., 2021) but also horizontally from an infected male to uninfected females during mating (Fiorenza *et al*., 2025; Son *et al*., 2021). Indeed, *s*Gff can be transferred to the female via the spermatophore, together with the ejaculate, and it can subsequently migrate to the spermathecae where spermatozoa are stored after mating (Fiorenza *et al*., 2025). In terms of tissue localization *s*Gff has been detected with particularly high densities in the testes, in male accessory glands, ovaries and in the hemolymph (Doudoumis *et al*., 2017). The presence of *s*Gff affects the mating behaviour of *Gff* (Fiorenza *et al*., 2025). It induces a significant bias in mating preferences, with mating pairs more frequently comprising individuals with the same infection status than would be expected under random mating conditions resembling the reinforcement process (Butlin & Smadja, 2018) described in other insect parasites, such as *Wolbachia* in infected tephritid flies (Jaenike *et al*., 2006). The reinforcement may reflect a form of adaptive coupling rather than a type of premating isolation. Whether this type of *s*Gff induced mating behavior contributes to the formation of the population structure of distinct host races is an open question (Bruzzese *et al*., 2022; Saarman *et al*., 2023)

On this background, a deeper understanding of *s*Gff influence on *Gff* reproductive biology is critical for evaluating its potential as a biological control agent. In this study. a transcriptomic approach and an autofluorescent assay have been used to determine the *s*Gff effect on the functionality of genes expressed in testes and MAGs. Such an approach offers an unprecedented molecular perspective on how this symbiont interacts with key reproductive processes such as fertility, immune response, seminal protein composition, sperm motility and functionality, mitochondria dynamics and energy metabolism.

## Materials and Methods

### *Gff* strain and rearing conditions

Flies used in this study were obtained from the *Gff* colony maintained at the Joint FAO/IAEA Center, Insect Pest Control Laboratory (IPCL) in Seibersdorf, Austria. This colony was established in 1986 using flies collected from the Bouar Provence, Central African Republic. Experimental flies were reared at 24 ± 0,5°C and 75 ± 5% relative humidity. Before adult eclosion, pupae were kept separated and as soon as adults emerged males were separated from females. Each newly emerged male was screened for *Spiroplasma* infection through careful surgical removal of one intermediate leg, performed under sterile conditions. Between one specimen and the following both forceps and scissors, used for the leg excision, were sterilized through brief immersion in 70% EtOH followed by exposure to the flame of a Bunsen burner. The excised leg of each specimen was collected in a new tube, and the respective fly was placed in an individual fly-holding cage. Then, the excised legs were used for *Spiroplasma* infection detection via PCR. Each male was maintained singularly, to avoid cross-contamination between individuals, until dissection on the sixth day. Adult flies were fed every two days for 10-15 min on defibrinated bovine blood using a silicone membrane (Bauer & Wetzel, 1976) and the first blood feeding occurred the day after eclosion, once the *s*Gff infection status was assessed to allow the separation of *s*Gff-infected flies from *s*Gff-uninfected flies during the blood feeding.

### Determination of the infection status

DNA extraction on the excised legs was performed using the ZR *Quick*-DNA 96 kit (Zymo Research, California, USA). A multiplex PCR using two specific primers pairs was used to assess the infection status of the adult males and verify the quality of the extracted DNA: Spi16s primers (F: 5’-GGGTGAGTAACACGTATCT-3’, R: 5’-CCTTCCTCTAGCTTACACTA-3’), a *Spiroplasma*-specific primer pair allowed the identification of the bacterial infection, and a *Gff* specific primer pair amplifying the housekeeping gene ß-tubulin (F: 5’-ACGTATTCATTTCCCTTTGG-3’, R: 5’-AATGGCTGTGGTGTTGGACAAC-3’) (Son *et al*., 2021). The PCR reaction was performed using the following cycling conditions: 94°C for 5 min, 34 cycles of 94°C for 45 sec, 58°C for 45 sec, 72°C for 1 min, and a final extension at 72°C for 10 min. Amplification products were electrophoresed on 2% agarose E-Gel (Fischer Scientific) stained with SYBR Safe.

### Dissections and RNAseq sample preparation

6-days-old virgin males were dissected under a stereomicroscope in sterile PBS 1X (Phosphate Buffer Saline), and for each male, testes and male accessory glands (MAGs) were collected; dissections were conducted carefully and all the fat body surrounding the reproductive tract was removed. For both *s*Gff-infected and *s*Gff-uninfected 6 pools of testes and 6 pools of MAGs were obtained, each pool containing tissues collected from ten males. Immediately after dissection, the dissected tissues were put into a sterile tube containing ice-cold 500 μl of TRIzol Reagent (Thermo Fischer Scientific) and, once the pools were complete, tissues were pestled and stored at -80°C.

### RNA extraction and library preparation

The resulting 24 samples were shipped to Edinburgh Genetics Ltd for RNA extraction and quality control (QC) assessment. All 24 samples passed the QC process. The library construction was carried out following Illumina standard protocols, and total RNA sequencing was performed.

### Differential gene expression analysis

The paired-end sequences from testes and MAGs of *s*Gff-infected and *s*Gff-uninfected tsetse flies were analysed through the nf-core/rnaseq pipeline version 3.12.0 (Di Tommaso *et al*., 2017; P. A. Ewels *et al*., 2020). Briefly, the pipeline consists of four steps: pre-processing, where raw reads are controlled and filtered; genome alignment and quantification; post-processing, where mapped reads are sorted, indexed, deduplicated and transcripts are assembled and quantified; finally, a multiQC file (P. Ewels *et al*., 2016) is produced, which summarizes results obtained from the pipeline. The STAR-Salmon method was employed, which uses STAR (Dobin *et al*., 2013) for the genome alignment and Salmon (Patro *et al*., 2017) for the expression quantification. The alignment was performed against the IAEA *Gff* annotated genome (Saarman *et al*., 2022) (version 63 of VectorBase, GCF_014805625.2 in NCBI). All the annotated proteins of the reference genome were blasted against the *D. melanogaster* database, wrapping BLASTP through NcbiblastpCommandline module from Biopython version 1.81 (Cock *et al*., 2009), to retrieve the most likely *D. melanogaster* ortholog gene symbols for each annotated protein.

The count matrix produced by the rnaseq pipeline was used to perform the gene expression analysis in R version 4.2.3 (R Core Team, 2022). Principal component analysis was performed using plotPCA from DESeq2 package (Love *et al*., 2014). Differentially expressed genes (DEGs) were detected employing the DESeq2 package (Love *et al*., 2014) and setting the log2fold change threshold to 1.5 and adjusted p-value threshold to 0.05. Afterwards, DEGs were employed for over-representation analysis (ORA) using the clusterProfiler package version 4.9.0 (Wu *et al.,* 2021) and *Glossina fuscipes fuscipes* gene ontology (GO) terms provided in the GAF file downloaded from NCBI (GCF_014805625.2_Yale_Gfus_2_genomic.gaf) integrated with the *D. melanogaster* GO terms for orthologous genes. biomaRt (Durinck *et al.,* 2005, 2009) was employed to ease the translation of gene symbols into gene identifiers. Plots were generated using ggplot2 (Wickham, 2011).

Metabolic pathways affected by *Spiroplasma* were detected using STRING (Szklarczyk *et al*., 2023). DEGs having a *D. melanogaster* ortholog were used for this analysis, as *D. melanogaster* served as the reference organism. Afterwards, connected subnetworks were selected, and metabolic pathways associated with them were identified by inspecting KEGG and Reactome enrichment results.

GO terms and metabolic pathways affected by *s*Gff were further investigated by performing a co-expression network analysis using the WGCNA package version 1.73, followed by ORA on subnetworks containing DEGs (Langfelder & Horvath, 2008; 2012).

### Microbiota analysis

After mapping the reads to the *Gff* genome, any unmapped reads were subsequently remapped to the *s*Gff genome (Bruzzese *et al*., 2025a) using the nf-core/rnaseq pipeline, with settings carefully tuned for bacterial genomes, as recommended in the user manual.

Metataxonomic profiling of the reads was performed using the nf-core/hgtseq pipeline (Carpanzano *et al*., 2022; Di Tommaso *et al*., 2017; P. A. Ewels *et al*., 2020). Briefly, unmapped reads were classified using Kraken2 (Lu *et al*., 2022; Wood *et al*., 2019; Wood & Salzberg, 2014), and the metataxonomic profiles of each sample were stored in a TSV file. The Kraken2 database was constructed by supplementing the standard database with all available reference genomes of *Mollicutes* (*s*Gff class; Table S1) and the genome of *Glossina fuscipes fuscipes*. The hgtseq pipeline also flagged likely contaminant taxa (Carpanzano *et al*., 2022), which were excluded from subsequent analyses. An additional decontamination step, based on a frequency approach, was carried out in R using the *decontam* package (Callahan, 2017/2025). Briefly, the likelihood of a taxon being a contaminant was estimated based on its frequency across samples. Finally, *phyloseq* and *DESeq2* (Love *et al*., 2014) were used to identify bacteria that were differentially present in relation to *Spiroplasma* infection.

### Autofluorescence assay on *Gff* sperm bundle

A mating experiment was performed using virgin *Gff* 8 days-old females and 6 days-old males. At the end of the copula, when individuals spontaneously separated, the spermatophore was immediately dissected from the female’s uterus, to avoid the dissolution of the spermatophore and the migration of the spermatozoa in the spermathecae (Attardo *et al*., 2020). Dissections were conducted under a Stereomicroscope (Leica MZ12) in sterile PBS 1X and the sperm bundle was isolated from the spermatophore capsule; for each mating pair, the legs of the males were collected for the determination of the infection status. The sperm bundle was transferred to a clean microscope glass slide, previously washed with a 50% alcohol-ether solution, and allowed to dry at room temperature. The spermatophore capsule was subjected to the same treatment. The microscope slides were then stored in the dark at room temperature until the autofluorescence assay. Twenty samples of sperm bundles and the relative spermatophore capsule were collected and analysed. Genomic DNA was extracted from the excised male legs using the protocol described in Baruffi *et al*. (1995) (Baruffi *et al*., 1995) and subsequently, a PCR with *Spiroplasma*-specific primers was performed, as described before. Autofluorescence (AF) spectroscopy was performed at IGM-CNR on unfixed, unstained sperms under epi-illumination, using a fluorescence microscope (Leitz, Wetzlar, Germany) integrated with a PMA 12 Optical Multichannel Analyzer (Hamamatsu Photonics Italia s.r.l., Arese, MI). The light excitation source was a LED-366 nm (Hamamatsu photonics) with two separate optical fibres: one to guide the excitation light to the samples, and another to collect the autofluorescence emission for the measuring system. Excitation and emission light have been selected through autothermal (KG1-BG38) and interferential (366 nm, T366 = 25%) filters, a dichroic mirror (390 nm, T366 < 2%) and a long pass filter (> nm). Spectra were collected in the 400–750 nm interval.

### Statistical analysis

R version 4.2.3^31^ was used to perform all statistical analyses. In order to assess the significance of comparisons concerning tissues and *s*Gff infection status it was employed the Mann-Whitney U test (or Wilcoxon signed-rank test)^44^ using wilcox_test from rstatix package^45^. In case of multiple comparison, Benjamini-Hochberg p-value correction was employed^46^.

## Results

### Expression profile of genes in *Gff* testes and MAGs

A total of 1.40 billion high-quality reads was obtained from the 24 biological samples (60.94 million reads per sample on average). No significant difference in the number of mapped reads between MAGs and testes (z = 24, n1 = n2 = 12, P-value = 0.266) and between positive and negative (z = 24, n1 = n2 = 12, P-value = 0.266) was observed. To have a picture of the gene expression profile of testes and MAGs, only *s*Gff-uninfected samples were considered, in order to avoid the influence of *s*Gff on gene expression of these two tissues. A total of 10,041 transcribed genes were identified among the twelve uninfected samples.

The principal component analysis (PCA) of Figure 4.1 shows a clear-cut division between the expression pattern of these two tissues. It appears clear that most of the gene expression variability is represented in the first principal component (PC1: 91%). Genes were then sorted by their mean expression level across the 12 *s*Gff-uninfected samples and clustering was performed based on the sample-wise gene expression levels of the first 50 genes, corroborating the clear division between MAGs and testes pointed out by the PCA.

**Fig. 1.**
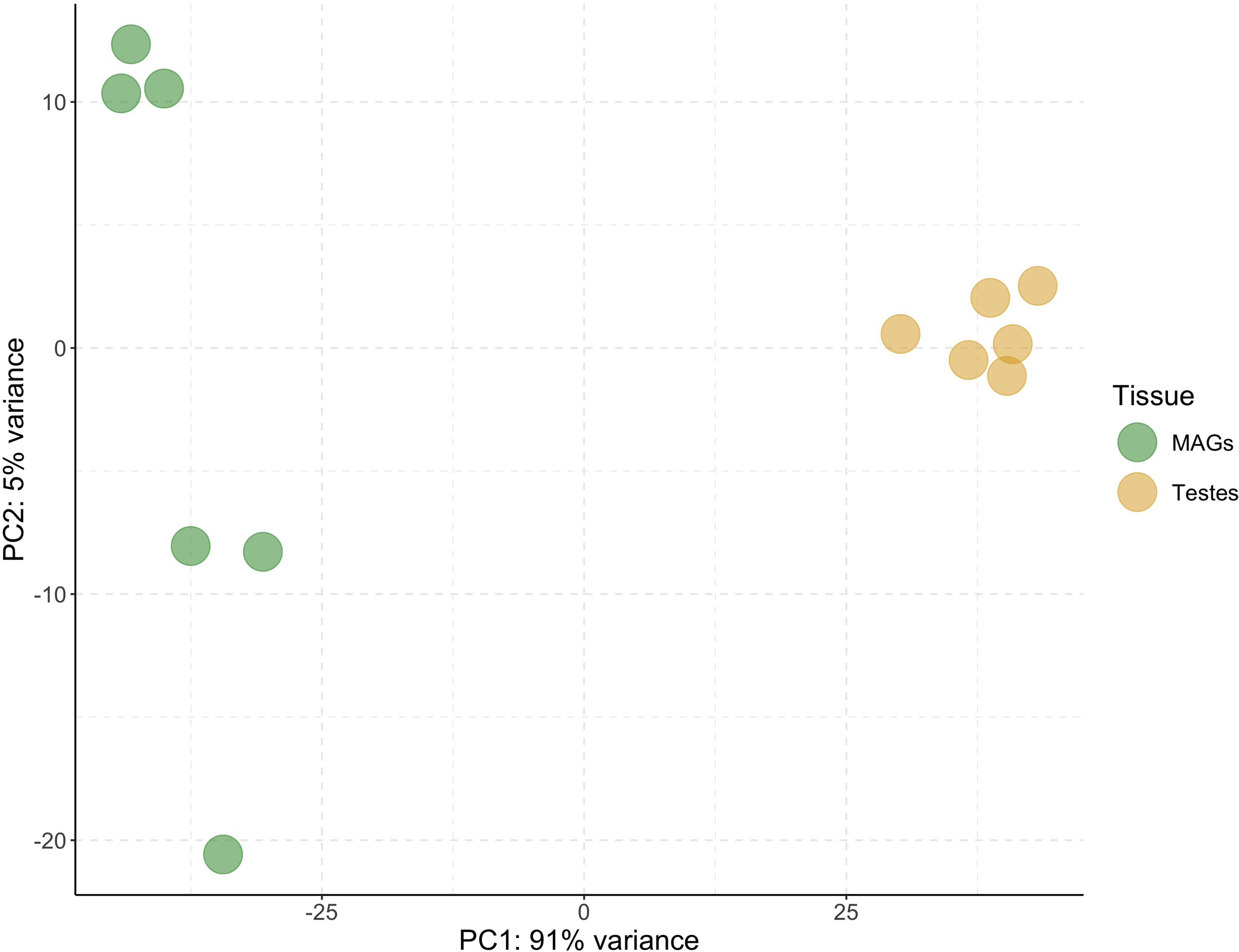
Principal component analysis (PCA) plot considering the first two PC calculated on the normalized counts of *s*Gff-uninfected samples. PC1 is the major component distinguishing the two tissues, which explains the 91% variance between testes and MAGs. MAGs = male accessory glands; PC = principal component

A heatmap of gene expression profiles in testes and MAGs has been derived (Figure 4.2). In this map, which considers the first 50 most expressed genes, the MAGs genes display a greater overall expression level than the testes ones. The data obtained from uninfected testes and MAGs was the basis for inferring the effect of *s*Gff on the expression profile these two tissues performed in infected flies.

**Fig. 2.**
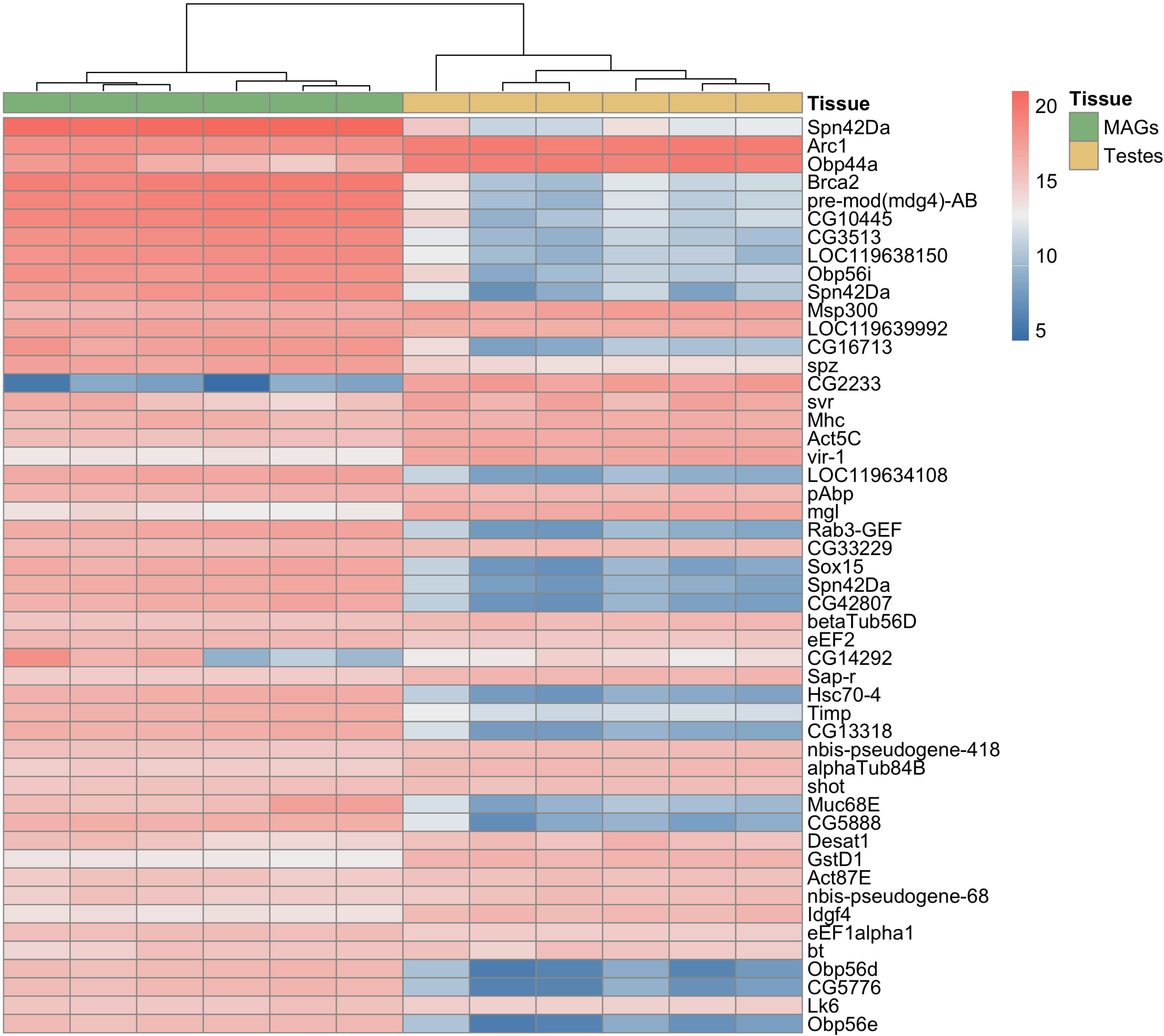
Heatmap of the 50 most highly expressed genes across uninfected samples. When available, *Drosophila melanogaster* orthologs are shown to the right of each row; otherwise, the original *Glossina fuscipes fuscipes* gene symbols are retained. Colors range from blue (lower expression) to red (higher expression) and represent sample-normalized counts for each gene. The heatmap highlights that MAGs are generally characterized by higher transcriptional activity compared to testes (see also Fig. 2a). MAGs = male accessory glands.

The following differential gene expression analysis (DGEA) between *s*Gff-uninfected MAGs and testes reported 1,173 differentially expressed genes (DEGs), setting the log2 fold change cut-off to 2 and -2 and the adjusted P-value cut-off to 0.05: 689 DEGs were upregulated in testes compared to MAGs and 484 downregulated in testes compared to MAGs (Figure 4.3a). The comparison of DEGs expression level between MAGs (mean = 6.09, sd = 2.17) and testes (mean = 5.31, sd = 2.04) showed a significant difference in expression between the two tissues (t_960_ = 6.23, *P* = 7.06e^-10^; Figure 4.3b), corroborating clustering hypothesis on MAGs-biased genes expression greater than testes-biased genes.

**Fig. 3.**
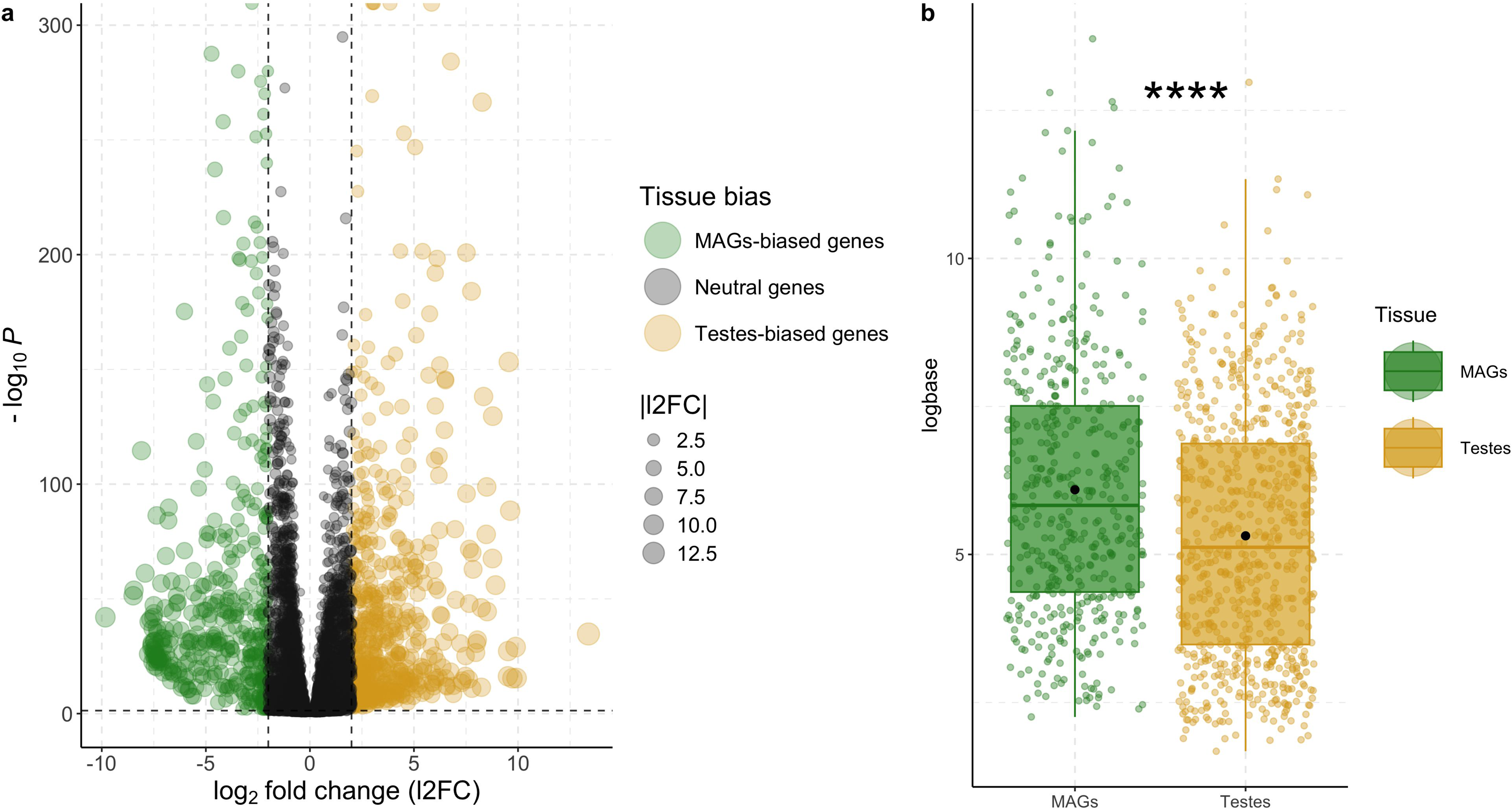
**(a)** Volcano plot showing differentially expressed genes of *s*Gff-uninfected samples (l2FC ≥ 2 and Padj ≤ 0.05) MAGs = male accessory glands. **(b)** Boxplot comparing the gene transcription (normalized read counts) of differentially expressed genes between tissues considering uninfected samples, which is significantly different (P-value = 7.06e^-10^; ****). The two tissues Black points define the mean of the two groups (MAGs = 6.09, testes = 5.31). Jitters show genes significantly differentially expressed and biased to the two tissues, which are more in testes (n = 739) compared to MAGs (n = 473). MAGs = male accessory glands.

An ORA using GO terms was performed to investigate the molecular functions (Figure 4.4) and biological processes (Figure 4.5) significantly impacted by DEGs. Twenty-two molecular functions and 32 biological processes were found significantly influenced by DEGs upregulated in testes, whereas 27 molecular functions and 35 biological processes were found significantly affected by MAGs-biased DEGs.

**Fig. 4.**
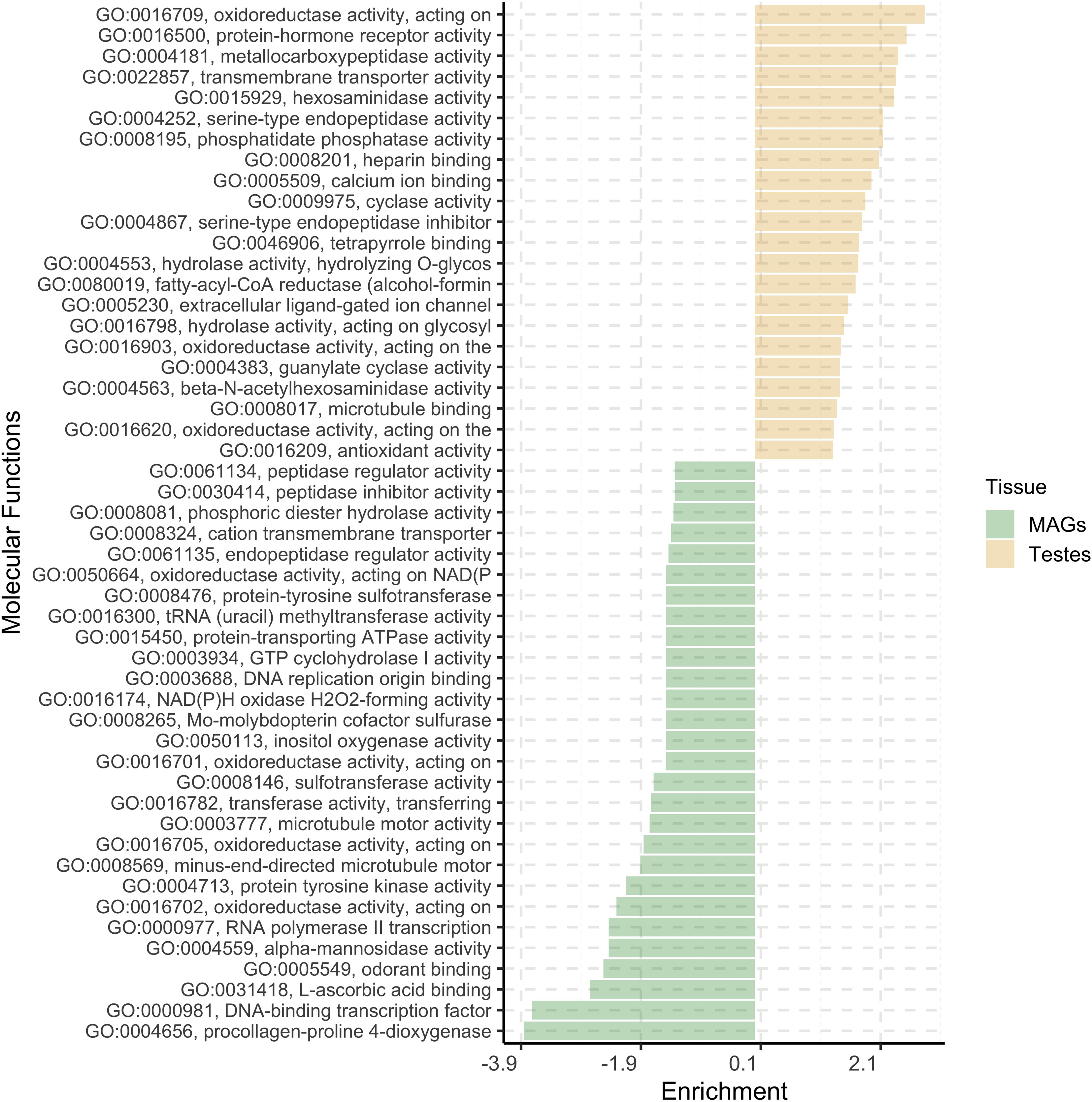
Gene set enrichment analysis results considering molecular function GO terms of *Glossina fuscipes fuscipes* genome annotations. x-axis reports Enrichment value, which is the elimination of the Kolmogorov-Smirnov (KS) statistics provided by topGO singed based on the gene set for which it was tested (positive values refer to testes-biased DEGs, negative values to MAGs-biased DEGs).

**Fig 5.**
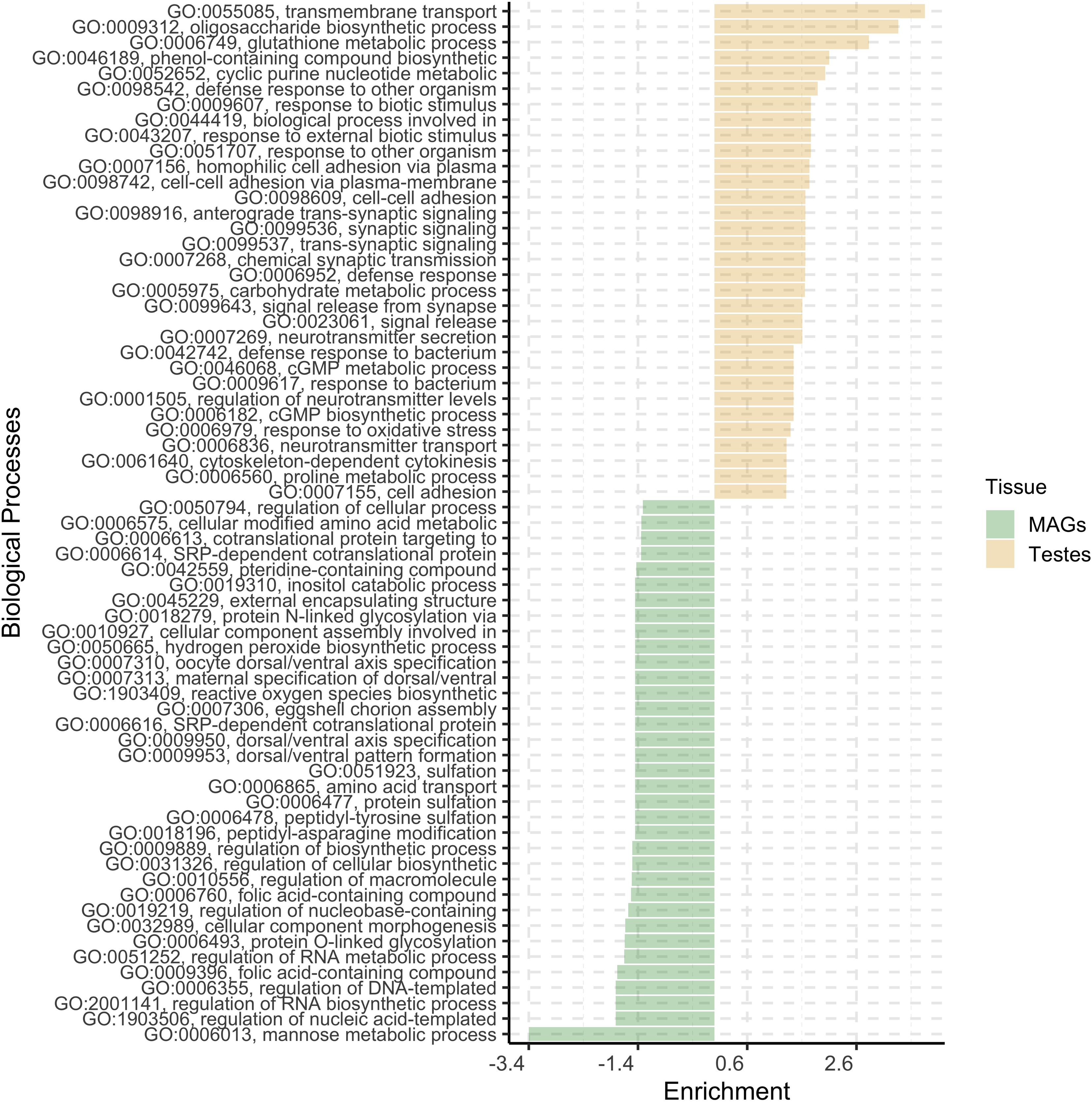
Gene set enrichment analysis results considering biological processes GO terms of *Glossina fuscipes fuscipes* genome annotations. x-axis reports Enrichment value, which is the elimination of the Kolmogorov-Smirnov (KS) statistics provided by topGO singed based on the gene set for which it was tested (positive values refer to testes-biased DEGs, negative values to MAGs-biased DEGs).

Regarding the biological process, the three most represented categories in testes were oxidoreductase activity, protein-hormone receptor activity and metallocarboxypeptidase activity, while in MAGs were procollagen-proline 4-dioxygenase and DNA-binding transcription factor. For the molecular function the most represented categories were transmembrane transport, oligosaccharide biosynthetic process and glutathione metabolic process in testes and the mannose metabolic process in MAGs.

### *s*Gff effect on gene expression in testes and MAGs

DGEA analyses were performed to assess the effects of *Spiroplasma* on MAGs and testes gene expression. A total of 9,408 transcribed genes were identified in the twelve MAGs samples. Only one DEG, corresponding to *Cipc Drosophila* ortholog (Figure 4.6), was significantly upregulated (P-value = 0.005, l2FC = 0.77) in the *s*Gff-infected MAGs, whereas no downregulated genes were identified.

**Fig. 6.**
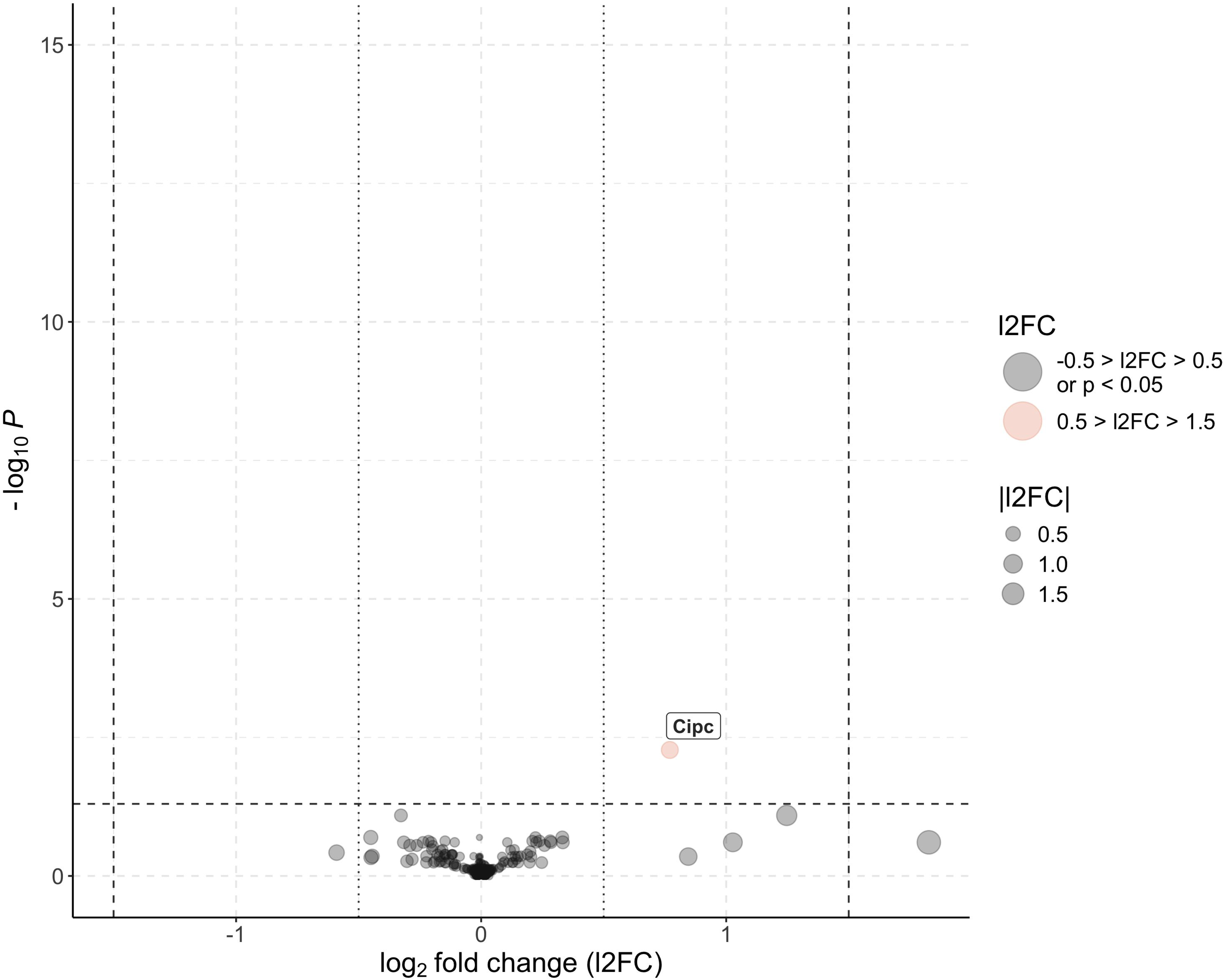
Volcano plot showing differentially expressed genes in *s*Gff-infected and *s*Gff-uninfected male accessory glands (MAGs). Genes with log= fold change (l2FC) > 0.5 and adjusted P-value < 0.05 (-log==P > 1.3) are significantly upregulated in *s*Gff-infected samples (shown in red), while those with l2FC < -0.5 and adjusted P-value < 0.05 are significantly upregulated in *s*Gff-uninfected samples. Genes with -0.5 < l2FC < 0.5 and adjusted P-value > 0.05 are not significantly differentially expressed (shown in black). Dashed black lines indicate the thresholds for significance (adjusted P = 0.05, -log==P = 1.3) and fold change (|l2FC| = 0.5). Point size corresponds to the magnitude of |l2FC|, and differentially expressed genes are labeled.

Conversely, 88 significant DEGs were identified between positive and negative testes out of 10,151 transcribed genes, considering a cutoff of l2FC = 0.5 (Figure 4.7). This roughly corresponds to a fold change of 1.42 minimum differential gene expression between positive and negative samples.

**Fig 7.**
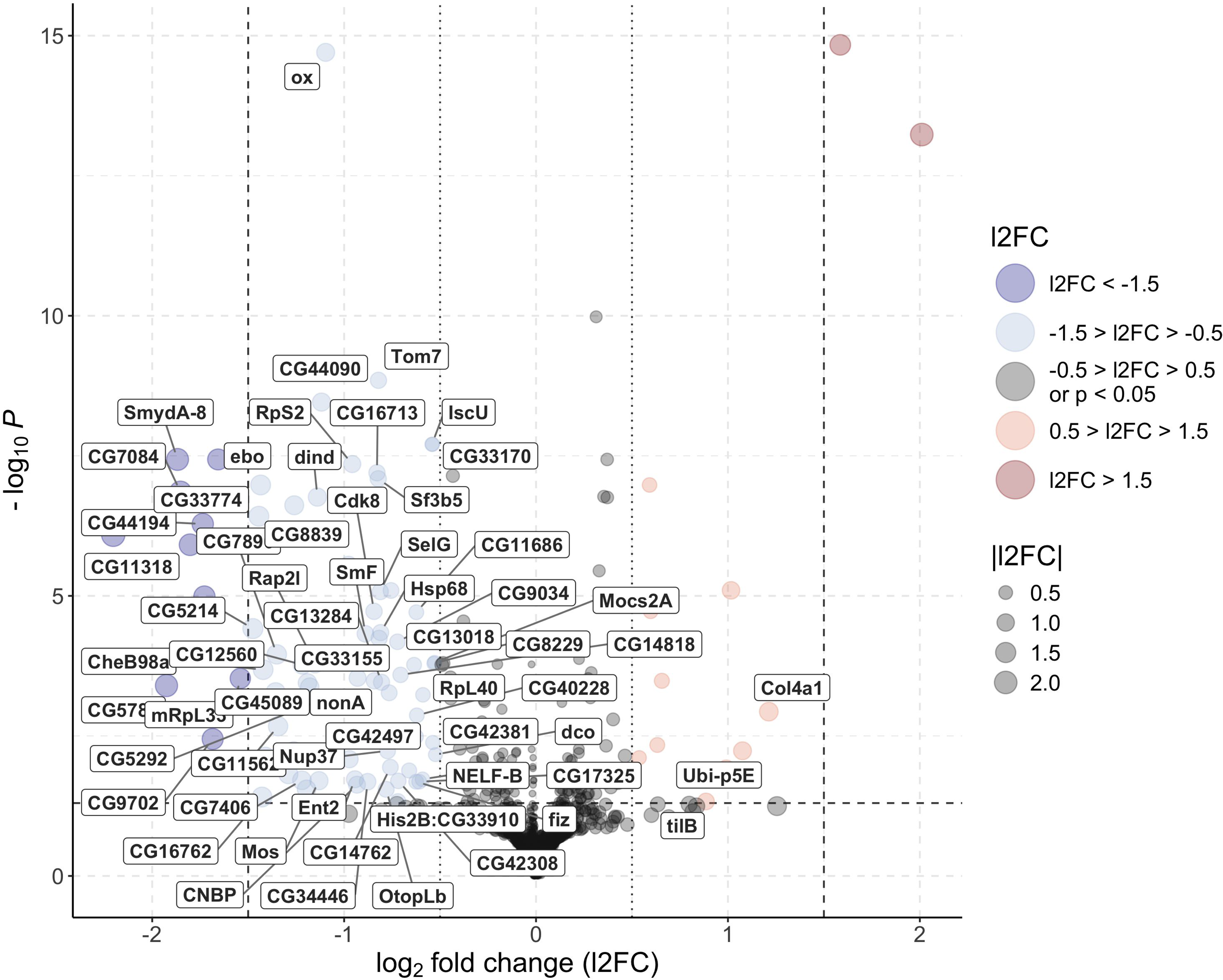
Volcano plot showing differentially expressed genes in *s*Gff-infected and *s*Gff-uninfected testes. Genes with l2FC > 1.5 and adjusted P-value < 0.05 are significantly upregulated in *s*Gff-infected samples (red), while those with l2FC > 0.5 and adjusted P-value < 0.05 are moderately upregulated (light red). Conversely, genes with l2FC < -1.5 and adjusted P-value < 0.05 are significantly upregulated in *s*Gff-uninfected samples (blue), while those with l2FC < -0.5 and adjusted P-value < 0.05 are moderately upregulated (light blue). Genes with -1.5 < l2FC < 1.5 and adjusted P-value >0.05 are not significantly differentially expressed (black). Dashed black lines indicate thresholds for significance (-log==P = 1.3, |l2FC| = 1.5), and dotted black lines indicate |l2FC| = 0.5. Point size corresponds to the magnitude of |l2FC|, and differentially expressed genes with *Drosophila melanogaster* orthologs are labeled.

Of the 88 DEGs (73 of which have a *D. melanogaster* ortholog), 66 (52 with *D. melanogaster* ortholog) were downregulated and only 10 (3 with *D. melanogaster* ortholog) upregulated in positive testes compared to negative testes. The following DEGs were noted: *ctenidin-3-like* (downregulated), antimicrobial; *tilB* (upregulated) and *CG7896* (downregulated), involved in flagellar function; *ox* (downregulated), central component of the cytochrome c oxidase; *Gld* (downregulated), coding for the glucose dehydrogenase; and *Ubi-p5e* (upregulated) and *RpL40* (downregulated) involved in ubiquitin activity.

### Biological processes affected by *s*Gff in *Gff* testes

An ORA analysis was performed on the GO terms of the 88 DEGs. This analysis identified five biological processes which were downregulated in positive testes: anion transport, inorganic anion transport, sulfate transport, sulfur compound transport and mitochondrial cytochrome c oxidase. A co-expression analysis of the 10,151 transcribed genes in testes was also performed. It revealed that 59 of the DEGs identified through the DGEA are part of the same co-expression network, characterized by 523 genes (Figure 4.8).

**Fig. 8.**
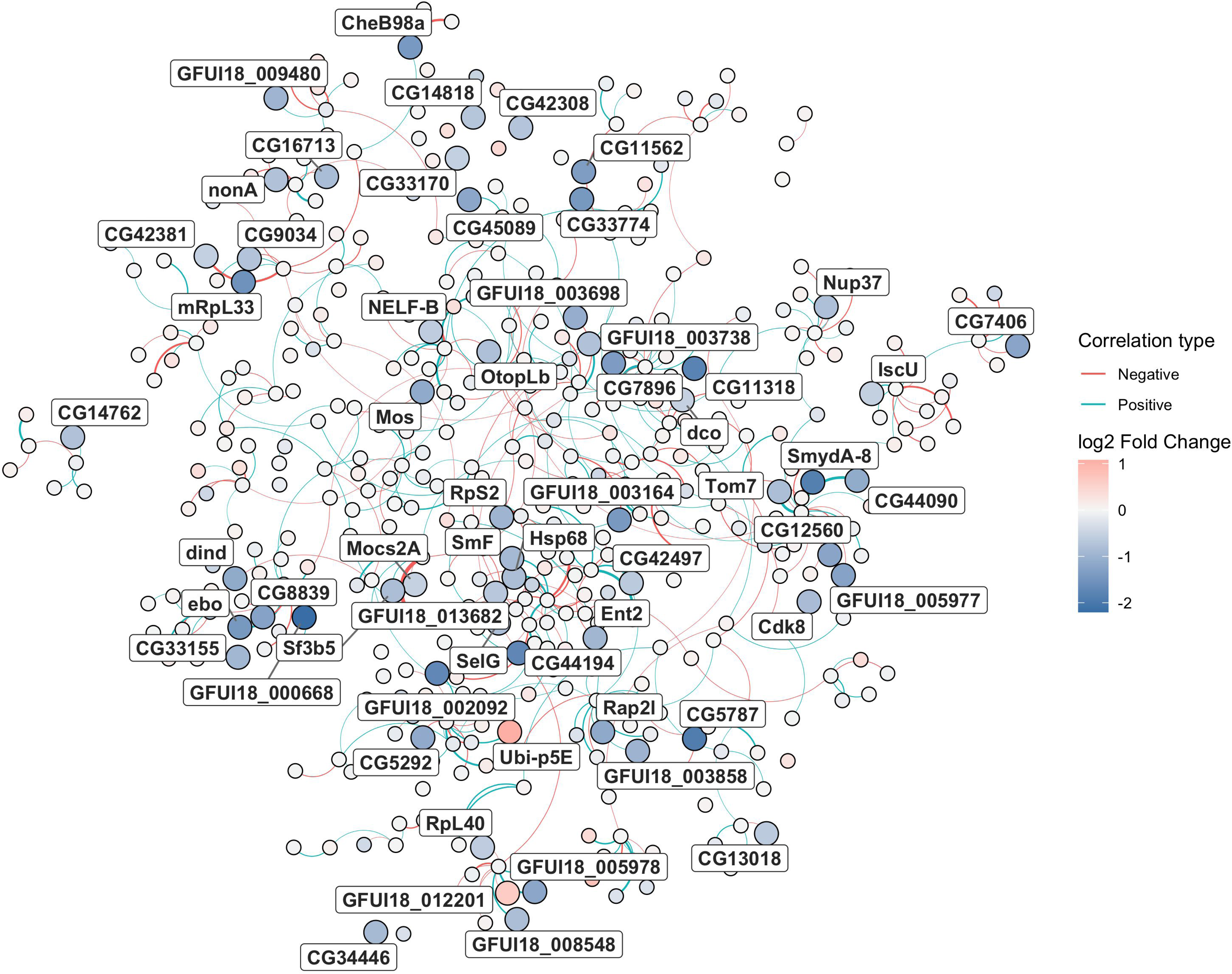
Network graph generated from the co-expression network analysis. Each point represents a gene that belong to the subset of genes strictly correlated with each other (523/10151 genes) and containing most of the differentially expressed genes (59/88 genes; reported in labels and greater node size). Nodes colors ranges from blue (downregulated in *s*Gff-infected testes) to red (upregulated in *s*Gff-uninfected testes) passing from white (neutral). Edges color represents the type of correlation (positive is red, negative is blue). The greater the correlation between nodes, the thicker the edges.

Subsequent ORA analyses showed that this co-expression network defines the overall mitochondrial activity including biological processes such as mitochondrial respiratory chain complex assembly, cytochrome complex assembly, cellular components such as mitochondrial respirasome, mitochondrial respiratory chain complex IV, ribonucleoprotein complex, ribosome and molecular functions such as structural constituent of ribosome.

The 73 DEGs having a *D. melanogaster* ortholog were submitted to STRING analysis, and they originated in two connected sub-networks based on GO and metabolic information (estimated and/or previously characterized according to the STRING database). One of the two networks was composed of three genes (Fig. 4.9b) and was associated with transcription elongation factor complex (CL:7140, STRING CLUSTER db). The other subnetwork was composed of 21 genes (Fig. 4.9a): three KEGG and 132 Reactome pathways were associated with it. Most of the Reactome results were due to the presence of *RpL40* and *Ubi-p5E* in the network, which are related to the ubiquitin activity.

**Fig. 9.**
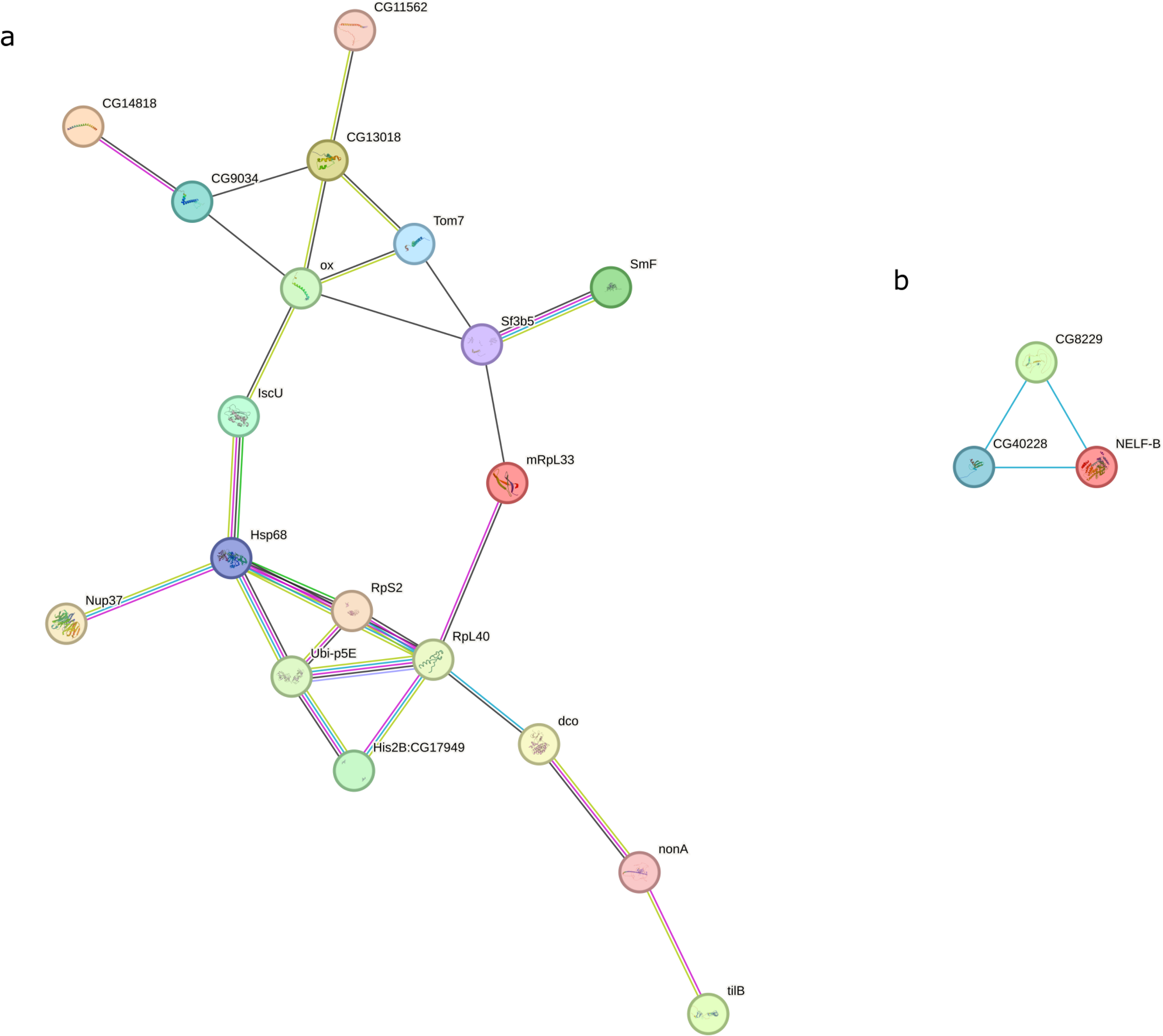
STRING results considering the 73 DEGs having a *D. melanogaster* ortholog. 21 of them where linked each other in a subnetwork. **(a)** A subnetwork of 21 interconnected DEGs. **(b)** A separate subnetwork of 3 DEGs. Nodes represent DEGs, while edges indicate interactions classified by color: blue for known interactions from curated databases, pink for experimentally determined interactions, light green for predicted gene neighbourhood associations, yellow-green for text-mined interactions, and black for predicted co-expression.

### *s*Gff impact on the sperm

To explore if *Spiroplasma* affects sperm function, the attention was centered on the sperm mitochondrial activity. To do this an assay which uses autofluorescence spectroscopy was used. The autofluorescence spectra collected from both spermatophores and the sperm bundle, isolated from spermatophore, covered in general the 400-600 nm spectral range, with a main band in the 400-550 nm region, ascribable to NAD(P)H emission. When normalized to 100 a.u. at the peak maximum, the emission spectral profiles from *s*Gff-infected or *s*Gff-uninfected spermatophores did not show remarkable differences. Compared to the spermatophores, sperm bundles showed, in general, a slightly higher tail in the yellow-red region (520-650 nm), and only *s*Gff-infected samples exhibited a shift of about 10 nm toward the blue region. The difference in autofluorescence real values between *s*Gff-infected and *s*Gff-uninfected sperm bundles resulted significant (*P* = 0,01; Figure 4.10; Table 1) for the Mann-Whitney U test.

**Fig. 10.**
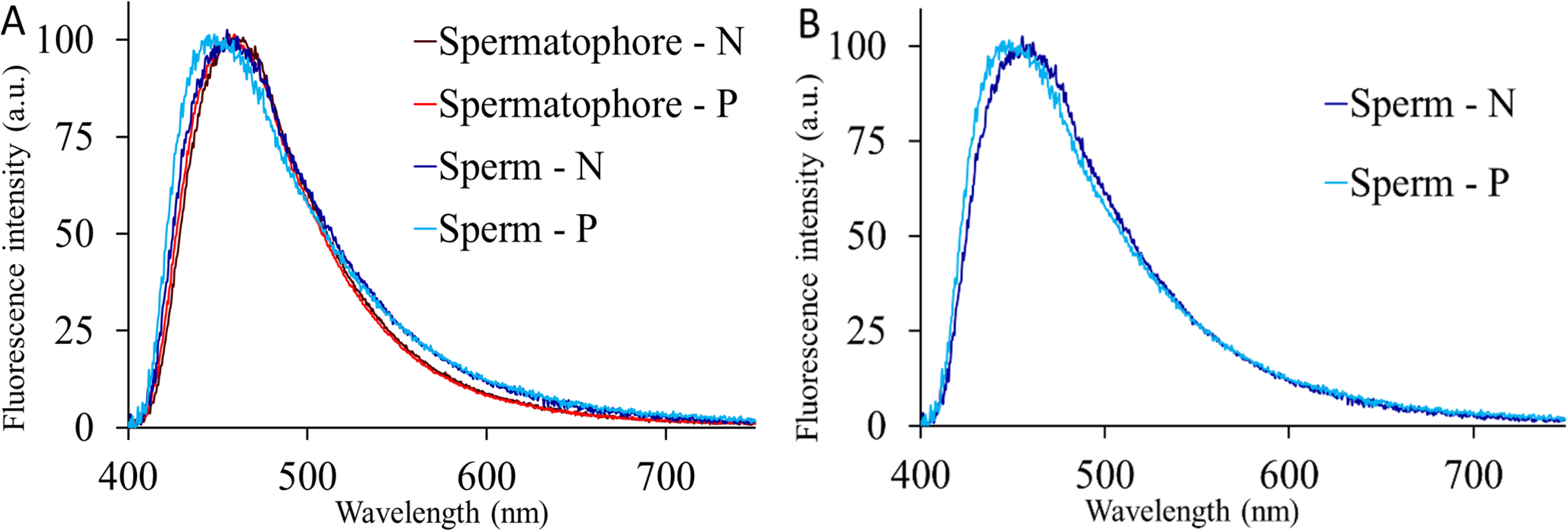
The curves represent the mean autofluorescence spectra from spermatophore and sperm bundle samples (A) collected from *s*Gff-uninfected (N) or *s*Gff-infected (P) individuals. (B) To better appreciate the slight blue shift of the emission from P compared to N spermatozoa, their emission spectra are shown also separately from those of the spermatophores. All spectra are normalized to the maximum peak value of 100 a.u. (fluorescence intensity) to facilitate the comparison of the emission profiles.

**Fig 11.**
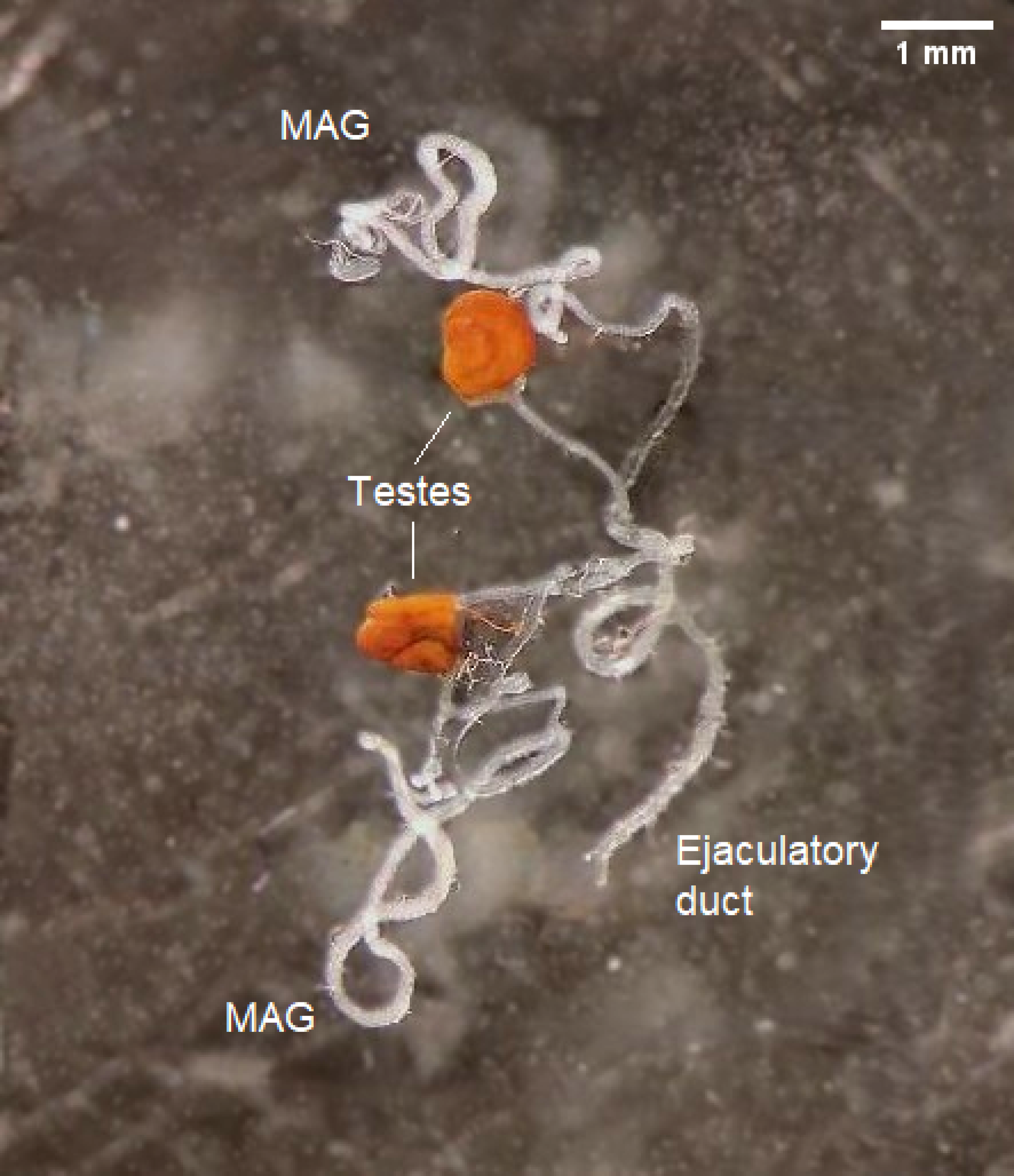
*Glossina fuscipes fuscipes* whole male reproductive tracts; MAG = Male accessory gland. Scale bar = 1mm.

**Fig 12.**
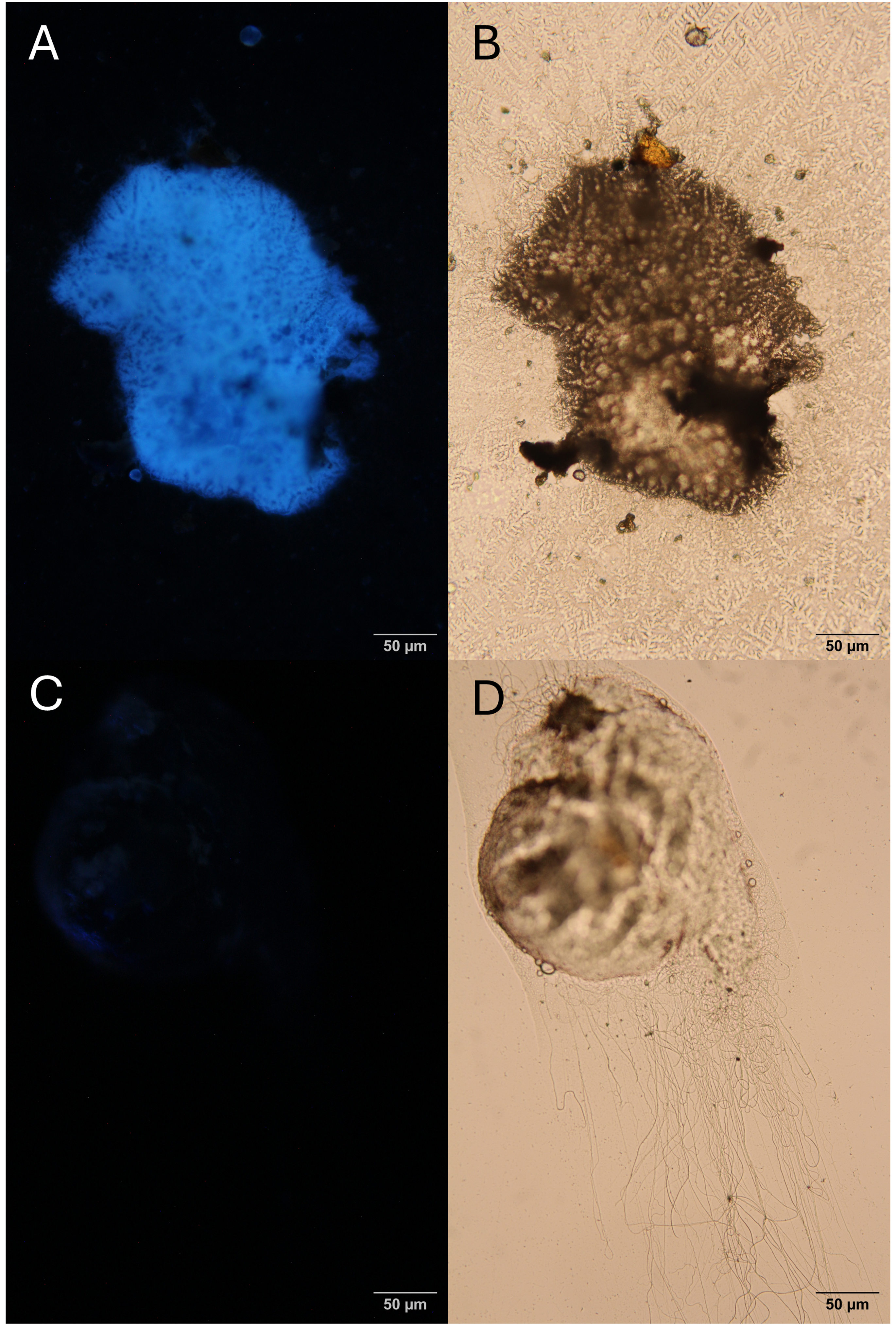
**(a)** *s*Gff-uninfected sperm bundle in fluorescent light, **(b)** *s*Gff-uninfected sperm bundle under microscope, **(c)** *s*Gff-infected sperm bundle and **(d)** *s*Gff-infected sperm bundle under microscope.

**Table 1.** Autofluorescence emission values measured from the spermatophore capsule and sperm bundle expressed as means (a.u.), Standard Deviations and Standard Errors. Data were calculated as integrated values of the 410-550 nm interval of each emission spectral area.

| Spermatophore capsule |  |  |  |  |
| --- | --- | --- | --- | --- |
|  | Mean | Std Dev | Std Error | Significance |
| Spermatophore N (n = 18) | 2184.11 | 1588.67 | 374.45 | NS |
| Spermatophore P (n = 35) | 1417.51 | 869.17 | 146.92 |  |
| Sperm bundle |  |  |  |  |
| Sperm bundle N (n = 16) | 1718.24 | 1450.25 | 362.56 | P < 0.05 |
| Sperm bundle P (n = 41) | 872.73 | 731.95 | 147.09 |  |

### *s*Gff gene expression in testes and MAGs dataset

To assess the potential presence of *Spiroplasma* transcripts in testes and MAGs, all the reads were mapped to the host genome searching for unmapped reads that could belong to the bacterium. It was found that the 93% of reads mapped to *Gff* genome (GCF_014805625.1), with a remaining proportion of unmapped reads (7%). These unmapped reads were mapped against *s*Gff genome (Bruzzese *et al*., 2025b). Among these unmapped reads, only four transcripts had a mean count of 10: ribosomal genes (23S and 16S rRNA), a transfer RNA (*<u>ssrA</u>*), and a ribonuclease (*RNaseP_bact_b*).

### *s*Gff and *Gff* microbiota interaction

To assess the *Gff* microbiota composition using the RNAseq data, a profiling approach through Kraken2 was applied to the testes and MAGs dataset. A total of 338 genera of bacteria in testes (mean abundance threshold across samples = 10) were identified. Considering bacteria genera having a minimum relative abundance of 0.1% in at least 3/12 samples, 59 genera resulted significantly differentially present in testes (Table 2). Among these, 49 genera had a significantly greater abundance in *s*Gff-infected, while 10 genera had a lower abundance. Among the most differentially abundant bacteria, in addition to *Spiroplasma* (l2FC = 2.89; baseMean = 30,616.42), were found *Wolbachia* (l2FC = 1.34; baseMean = 438.73) and *Wigglesworthia* (l2FC = 4.33; baseMean = 15,722.29). In MAGs, 263 genera were detected (mean abundance threshold across samples = 10), and no significant differences in genera abundance were found between infected and uninfected samples.

**Table 2.**
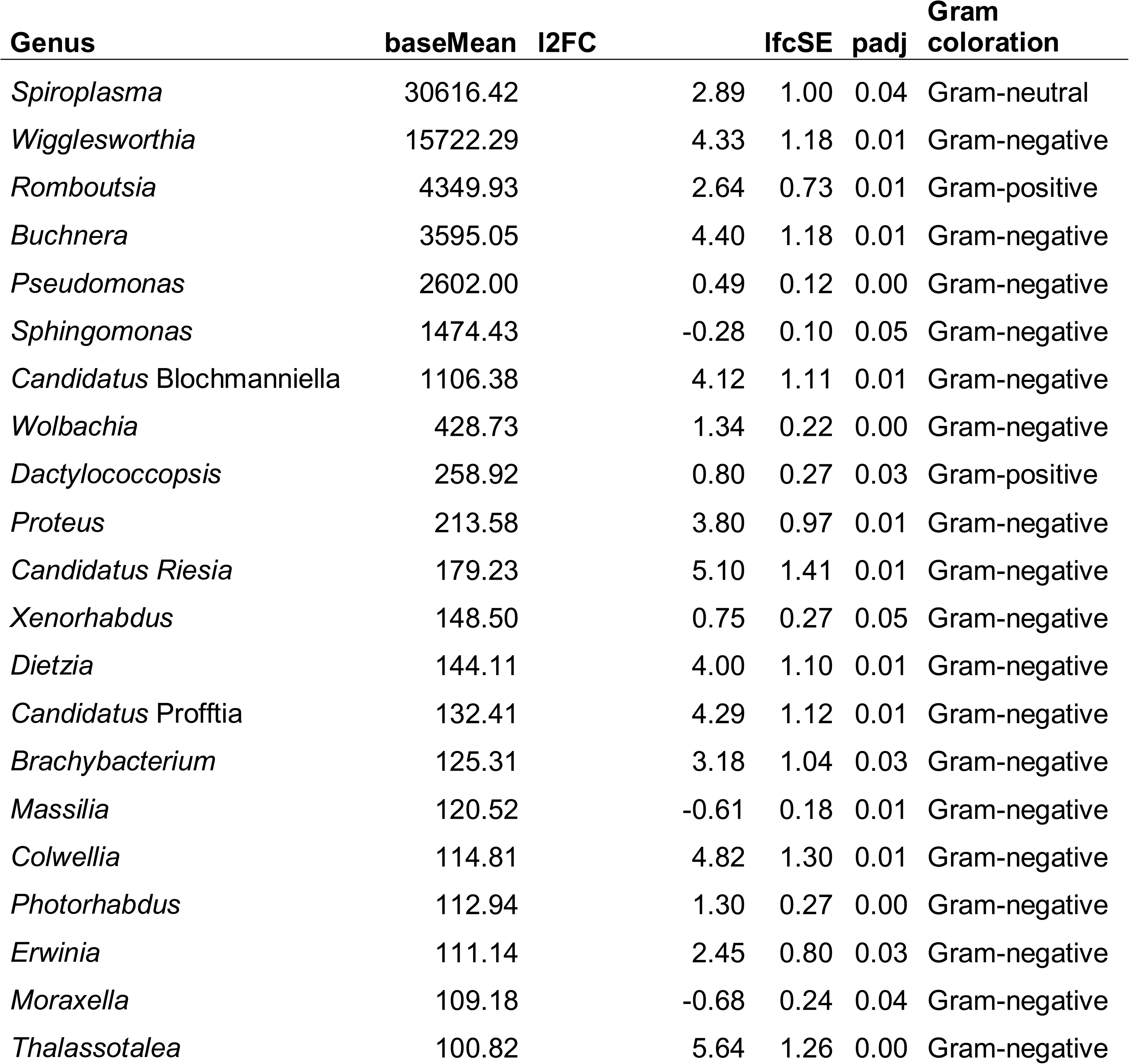
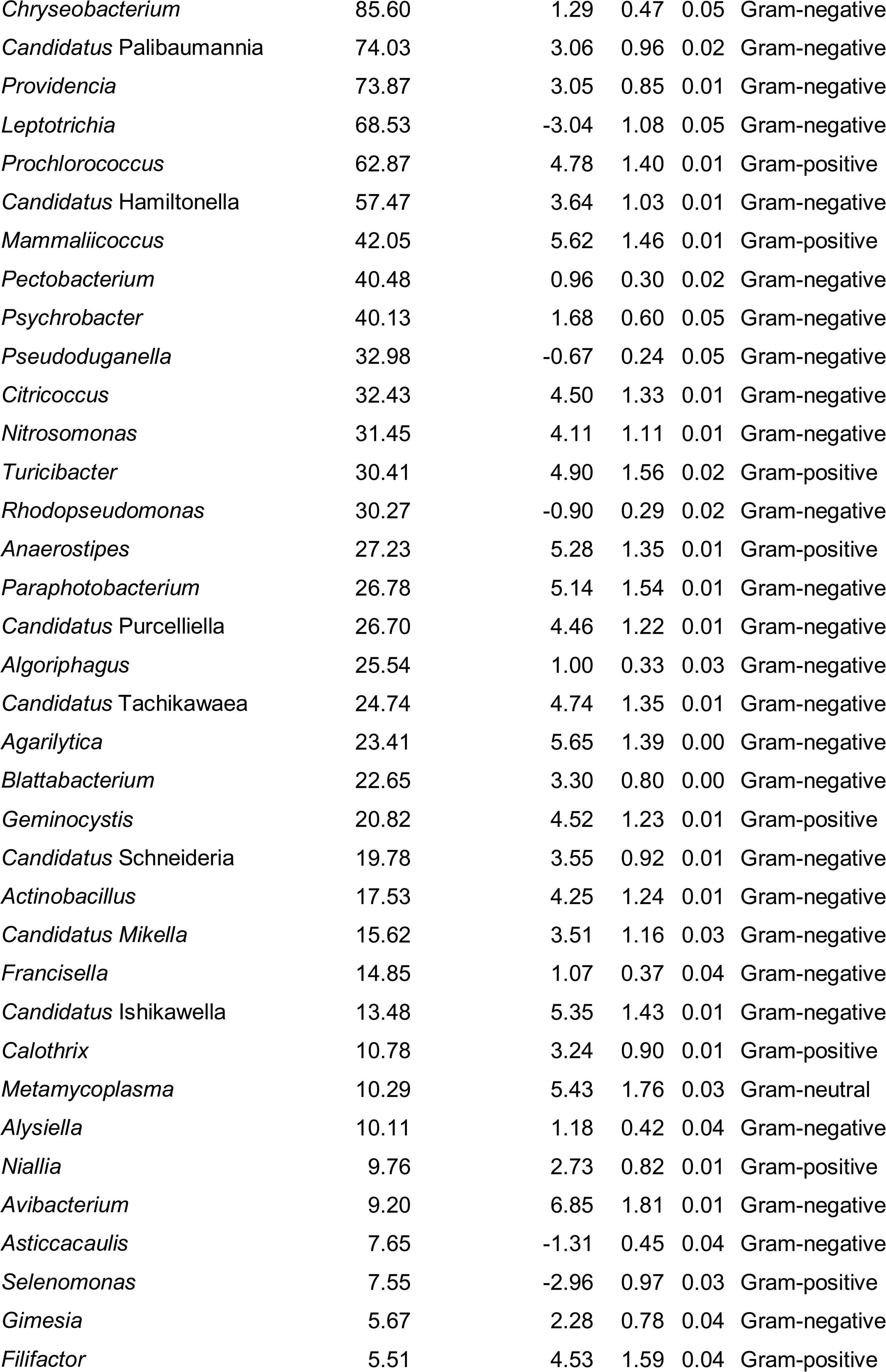
List of the differentially abundant bacteria found in the transcriptomic dataset.

## Discussion

### Gene expression profiles in testes and MAGs in the riverine species *Gff*

As Fig. 4.2 and Fig. 4.3 portray, when comparing MAGs and testes a lot of genes resulted differentially expressed (973 DEGs). While a greater number of genes is transcribed in testes (689 DEGs with l2FC higher in testes, 439 in MAGs; Fig. 4.3), the overall gene expression is greater in MAGs (5.31 logBaseMean in testes, 6.09 in MAGs; Fig. 4.3b), highlighting the differential but complementary role of these organs. These results were expected, as in male reproduction of other *Glossina* species, such as *G. morsitans*, where MAGs display a small number of highly expressed genes while testes contribution for sperm composition consists in a diverse and rich array of less expressed genes (Scolari et al., 2016; Attardo et al., 2020; Savini et al., 2021). While some of the DEGs upregulated in MAGs code for unknown functions, others code for enzyme inhibitors and peptidase regulators (Scolari et al., 2016; Attardo et al., 2020; Savini et al., 2021). On the other hand, DEGs upregulated in testes are associated with binding, oxidoreductase/transferase activities, cytoskeletal and lipid/carbohydrate transporter functions and mitochondrial functions. As previously shown (Scolari *et al*., 2016; Savini *et al*., 2021) sperm-associated genes are typically more conserved among the different *Glossina* species and insect taxa than the MAG ones. The high sequence conservation observed for sperm-associated genes may reflect essential sperm-specific processes they participate in. In contrast, MAGs genes code for key factors in species-specific male reproductive success and as such are subject to rapid evolution resulting from sexual conflict and competition that are very high in *Gff* (Savini *et al*., 2021). Indeed, in this riverine species from the Palpalis group, the selective pressure level in MAGs is higher than in savannah flies such as the *Morsitans* subgroup. *Gff* flies typically live in narrow riverine habitats and suffer seasonal demographic fluctuations. During the dry season, populations undergo a dramatic demographic contraction with the remaining flies concentrating in most refugia. At the end of the dry season, within the residual population emerging after the bottleneck, the strength of male competition increases because of the greater number of interactions for achieving copulation.

### *Spiroplasma* differentially affects testes and MAGs

*Spiroplasma* seems to marginally affect MAGs expression profile as only one DEG was detected when comparing infected and uninfected tissues. On the other hand, it deeply affects the gene expression profile of testes, with 88 DEGs detected when performing the same comparison. Assuming the *Gff* orthologs have a similar function associated with the *D. melanogaster* or *G. morsitans* counterparts, such genes characterizing testes affected several functions and metabolic pathways.

### *Spiroplasma* alters the circadian rhythm of reproductive tissues

Analysis of DEGs between sGff-infected and sGff-uninfected tsetse flies revealed a single upregulated gene (l2FC = 0.77) in MAGs. This transcript, the ortholog of *Cipc*, encodes a protein with a conserved repressive function within the CLK-CYC transcriptional feedback loop, a key regulator of the circadian rhythm (Rivas *et al*., 2021). In *D. melanogaster*, upregulation of *Cipc* has been associated with lengthening of the circadian period (Rivas *et al*., 2021).

In the testes, the gene *dco* was downregulated (l2FC = -0.52) in response to *s*Gff infection. The *dco* gene encodes Doubletime (DBT), a kinase crucial for circadian rhythm regulation. DBT phosphorylates PER (Period), targeting it for degradation. However, when cytoplasmic levels of TIM (Timeless) are high, PER binds to TIM, forming a complex that represses the CLK-CYC system upon DBT binding (Zheng *et al*., 2014). Previous studies have shown that DBT reduction lengthens the circadian rhythm due to desynchronization of the clock network, reduced PER phosphorylation, and accumulation of nuclear PER (Zheng *et al*., 2014). These findings suggest a potential slowing of the circadian rhythm in both MAGs and testes. This could consequently lead to disruptions in sperm production, fertility, and mating behavior.

It has been previously shown that circadian genes play a crucial role in reproductive fitness (Beaver *et al*., 2002). This same study indicated that *per* and *tim* mutants in *D. melanogaster* exhibited reduced sperm production and altered sperm quality, as well as impaired maturation and release from the testes, ultimately resulting in reduced fertility.

Additionally, the hypothesis of a mating behavior alteration is supported by the downregulation induced on *nonA* (l2Fc = -0.76), a sex-linked gene encoding an RNA-binding protein involved in visual behavior and courtship song of *D. melanogaster*. Mutation of *nonA* can disrupt the courtship song of flies (Rendahl *et al*., 1996). Another gene that was shown affecting the song behaviour in *Drosophila* was found upregulated: LOC119644721 (ortholog of *D. melanogaster* gene *tilB*; l2FC = 0.88) (Tauber & Eberl, 2001). The differential expression of both circadian rhythm and courtship genes may explain the altered mating behavior previously observed in *Gff* in relation to *s*Gff infection (Dera *et al*., 2025; Fiorenza *et al*., 2025). Specifically, infected flies exhibited better mating fitness compared to their uninfected counterparts, potentially due to the slowed circadian rhythm and altered courtship behavior caused by the differentially expressed genes in response to *s*Gff infection.

### *Spiroplasma* has a negative impact on the sperm viability and energy metabolism in testes

*s*Gff affects genes involved in flagellar functions. Indeed, the *D. melanogaster* ortholog gene *tilB*, upregulated in testes (l2FC = 0.88), not only is involved in courtship as described previously, but also affects flagellar function. Indeed, this gene encodes a protein essential for the transport and/or assembly of the cilium/flagellum, and alterations in its expression can lead to sperm flagella dysfunction in *D. melanogaster* (Kavlie *et al*., 2010). Conversely, LOC119640098, ortholog of *D. melanogaster* gene *atk*, was downregulated (l2FC = -1.35). *atk*, encodes a protein that surrounds the cilium and is critical for its assembly, structural integrity, chemosensation, proprioception (Andrés *et al*., 2014), and it has also been positively associated with sperm viability in *Apis mellifera* (McAfee *et al*., 2021). The differential regulation by *s*Gff – upregulation of *tilB* and downregulation of *atk* – presents a paradox. As *tilB* is primarily linked to flagellar assembly, while *atk* has broader roles, including chemosensation, proprioception, and flagellar integrity, the disruption of *atk* expression likely outweighs the benefit of *tilB* upregulation, leading to a net negative effect on sperm function.

Genes which display orthology with *D. melanogaster* orthologs associated with energy metabolism were also affected. For example, *Gld* (l2FC = -0.62) encodes glucose 1-dehydrogenase, a key enzyme in an alternative glucose metabolic pathway. This enzyme oxidizes glucose into gluconate, which feeds into the pentose phosphate pathway and generates NADPH, a key electron transfer molecule in various processes. Additionally, *CG7084* (l2FC = -1.85), which encodes a transmembrane cation transporter orthologous to the human solute carrier family 22 members 1-5 (SLC22A1-5), was downregulated. Notably, SLC22A5 encodes the organic cation/carnitine transporter 2, which in human and mouse is the main transporter involved in carnitine intake in reproductive organs, and mutations in this gene were linked to male infertility (Linn *et al*., 2021). Carnitine plays a central role in energy metabolism by shuttling long-chain fatty acids into mitochondria for β-oxidation, which produces ATP. Reduced carnitine transport could impair β-oxidation, leading to decreased energy availability for spermatogenesis and sperm function.

Mitochondrial energy production appears to be significantly disrupted by *s*Gff infection. Several genes critical to the citric acid (TCA) cycle and the electron transport chain (ETC) were downregulated. For instance, *CG5214* (l2FC = -1.47) encodes dihydrolipoamide S-succinyltransferase, part of the 2-oxoglutarate dehydrogenase complex, which catalyses the conversion of 2-oxoglutarate into succinyl-CoA in the TCA cycle (Araújo *et al*., 2013). Similarly, multiple genes involved in the ETC were downregulated, as reported by GSEA. *ox* (l2FC = -1.10) encodes ubiquinol-cytochrome c reductase subunit 9, a core component of Complex III of the ETC (cytochrome c reductase). Another gene, *CG42308* (or *sloth2*), component of Complex III, was found downregulated (l2FC = -0.72). This enzyme facilitates electron transfer and proton translocation, playing a central role in oxidative phosphorylation and ATP production (Frolov *et al*., 2000). *CG9034* (l2FC = -0.72) encodes NADH-ubiquinone oxidoreductase subunit A3, a key part of Complex I of the ETC (NADH dehydrogenase) (Agip *et al*., 2023; Pueyo *et al*., 2023). Additionally, genes involved in the assembly of cytochrome c oxidase (Complex IV of the ETC), including *CG44194* (l2FC = -1.73), *CG11562* (l2FC = -1.34), *CG13018* (l2FC = -0.64) and *CG33170* (l2FC = -0.54) were also downregulated. Complex IV is essential for oxidative phosphorylation, and its proper function is crucial for spermatogenesis, sperm motility, and viability (Y.-J. Park & Pang, 2021).

Mitophagy-related genes were also impacted, suggesting that *s*Gff infection disrupts mitochondrial homeostasis. *Tom7* (l2FC = -0.82) encodes translocase of outer membrane 7, a complex responsible for importing proteins into mitochondria. This protein also regulates mitophagy, the selective degradation of mitochondria, and mitochondrial fission (Maruszczak *et al*., 2022; Sekine *et al*., 2019; Wang *et al*., 2023). The downregulation of *Tom7* may impair mitochondrial quality control. In contrast, *Ubi-p5E* (l2FC = 1.07), encoding ubiquitin-5E, and *RpL40* (l2FC = -0.59), encoding a ribosomal 60S component fused with ubiquitin, are both part of the ubiquitin-proteasome system (UPS) (Zhang & Zhang, 2019), which is involved in mitophagy and the removal of damaged proteins (Chen *et al*., 2019; Y. Li *et al*., 2022; G. H. Park *et al*., 2021).

The cumulative effects of *s*Gff on genes associated with sperm flagella and energy metabolism point to a potentially harmful impact on testes function and, in turn, on sperms. One plausible explanation is that *Spiroplasma* competes for essential energetic substrates, thereby reducing their availability to testes cells (Paredes *et al*., 2016; Son *et al*., 2021). This competition may diminish mitochondrial activity and energy production (Son *et al*., 2021), ultimately impairing spermatogenesis and sperm quality by disrupting the expression of critical genes. This hypothesis is supported by the autofluorescence assay results from this study, which showed that sperm from *s*Gff-infected *Gff* had lower levels of NADH. This indicates a reduced energy availability and a lower mitochondria functionality in *s*Gff-infected sperms (Walker & Tian, 2018).

These findings align with a previous study (Son *et al.,* 2021) that reported decreased beat frequency in sperm from *s*Gff-infected *Gff*, suggesting reduced motility. This reduction in motility is likely to impact sperm competitiveness within the spermatophore. Together, the negative impact on the expression of genes involved in energy metabolism, the reduced NADH levels and decreased beat frequency observed in these studies strongly suggest that *s*Gff infection limits the energy available to sperm, leading to impaired function and competitiveness. Furthermore, this may be one of the factors contributing to the failure to establish 100% infected colonies in the laboratory (Dera *et al*., 2025).

### *Spiroplasma* facilitates colonialization of other bacterial species in testes

The downregulation of LOC119637240 (l2FC = -0.98; ortholog of GMOY006519-RA and identified as *ctenidin-3-like*) suggests that *s*Gff can alter the antimicrobial action in testes (Scolari *et al*., 2016). Ctenidins are a class of antimicrobial glycine-rich peptides that act in the innate immune response against bacteria. These antimicrobial proteins (AMP) are characterized by a signal peptide followed by the mature peptide, and they are likely to act in hemocytes or be released into the hemolymph (Baumann *et al*., 2010; S. Li *et al*., 2021). In particular, ctenidin-3 was associated with bacteriostatic actions against Gram-negative bacteria and reduced the growth rate of Gram-positive by 30% (Baumann *et al*., 2010).

We attempted to perform a metataxonomic analysis on our samples, although they were not specifically prepared for this purpose: no rRNA depletion steps were performed prior to sequencing, nor was sequencing deep enough to better recognize bacteria transcriptomes (Wahl *et al*., 2022). Indeed, when aligning unmapped reads (with respect to the *Gff* genome) to the *s*Gff genome, we were only able to retrieve rRNA genes. This is because, in prokaryotes, the amount of rRNA is significantly higher than that of mRNA (Rosenow *et al*., 2001; Rosset *et al*., 1966; VanBogelen & Neidhardt, 1990; Wahl *et al*., 2022). Without rRNA depletion or deep sequencing, the imbalance in concentrations makes it extremely difficult to retrieve bacterial mRNA (Wahl *et al*., 2022). Additionally, it is important to note that only known bacterial species with documented interactions or associations with *Gff* were included in the Kraken2 database for profiling the samples from a metataxonomic perspective. This inevitably excludes bacterial species that were present in the samples but not considered due to database limitations (Liu *et al*., 2024). Moreover, the decontamination step also had limitations, as negative controls were unavailable to remove contaminants based on a “presence-based” approach (Callahan, 2017/2025). Instead, only contaminants flagged by the hgtseq (Carpanzano *et al*., 2022) or identified by decontam (Callahan, 2017/2025) through the ‘frequency-based’ approach were removed. However, our metataxonomic analysis provided valuable insights. It revealed that, in *s*Gff-infected testes, 17 bacterial genera increased in abundance, while only 7 genera showed a decrease. Notably, *s*Gff had the highest mean abundance across all samples (baseMean > 30k), confirming that *s*Gff is the primary bacterium distinguishing infected and uninfected samples, with no other bacteria showing comparable abundance. This suggests that the effect of *sGff* on *ctenidin-3-like* expression may be linked to changes in the abundance of other bacterial species in the testes.

One hypothesis is that *s*Gff competes with other bacterial species. As a wall-deficient bacterium, *s*Gff does not require the antimicrobial activity of *ctenidin-3-like*, potentially reducing the need for its production and resulting in its downregulation. However, our findings showed no strong impact on the abundance of bacterial genera that decreased in response to *s*Gff. This opens the door to another hypothesis: the high prevalence of *s*Gff may mask the presence of other bacterial species. This masking effect could prevent the immune system from recognizing Gram-positive and Gram-negative bacteria, thereby reducing the production of antimicrobials like *ctenidin-3-like*. As a result, less-dominant Gram-positive and particularly Gram-negative bacteria may proliferate in *s*Gff-infected testes.

Interestingly, among the 17 bacterial genera with increased abundance in *s*Gff-infected testes, we identified *Wolbachia*. *Wolbachia*, an intracellular parasite found in most arthropods (Alam *et al*., 2011; Werren *et al*., 2008), including *Gff* (Symula *et al*., 2013), is notable for its potential use as a biological control agent against disease vectors due to its ability to manipulate host reproductive fitness (Minwuyelet *et al*., 2023). Our results show that the quantity of *Wolbachia* in our samples is low, as shown by a previous study (Symula *et al*., 2013), but significantly different between infected and uninfected testes. It remains unclear whether ctenidin-3-like products act on intracellular parasites like *Wolbachia*, so we cannot conclude that the increased abundance of *Wolbachia* is linked to reduced ctenidin-3-like activity. Moreover, it is not clear whether *Wolbachia* can manipulate host reproductive fitness (Symula *et al*., 2013).

### Conclusion

Our study mainly provides evidence of testicular impairments in *s*Gff-infected males. Indeed, at the transcriptional level, *Spiroplasma* affects genes involved in mitochondria dynamics and energy metabolism, circadian rhythm, flagellar functions and sperm motility. This suggests that this bacterium may negatively affect male fitness in reproduction. From these data, two biological questions arise. The first is related to the potential use of *Spiroplasma* as a control method for the natural population of *Gff.* The second important question, strictly related to the first, pertains to the uneven distribution of *Spiroplasma* within the natural populations. Indeed, from preliminary studies it has been found that *Spiroplasma* may be absent in some individuals or vary in titers among different individuals in the population. In this context, we must also consider that our data was derived from a laboratory line that is maintained in stable condition. Knowing that natural populations of *Glossina fuscipes fuscipes* are subjected to seasonal fluctuation, further analyses are needed to verify if the presence of *Spiroplasma* and its effect on its host are constrained by this seasonal variation (Schneider et al., 2019).

## Author Contributions

R.P., G.F., A.M.M.A., F.F., F.L., G.G., S.A. and A.R.M. conceived and designed research; G.F., F.G., C.J.d.B. and K.M.D. collected samples and performed RNA library preparation; G.F. and A.C.C. performed the autofluorescence assay and imaging; F.L., R.P., and S.C. analyzed data; R.P., G.F., M.S., G.G. and A.R.M. interpreted results; R.P. drafted the manuscript with major input from G.F, G.G. and A.R.M.; A.R.M. has primary responsibility for final content. All authors read and approved the final manuscript.

## Acknowledgement

The authors would like to acknowledge technicians at the IAEA-Insect Pest Control Laboratory in Seibersdorf, Austria, for providing tsetse flies specimens used in this study, and PASS-Bio Med, Centro Grandi Strumenti of the University of Pavia, Pavia, Italy for the imaging of our samples with the transmission electron microscope (TEM).

