## Supplementary material for "Spiroplasma impairs testes gene expression in Glossina fuscipes fuscipes": Table S1

**Table S1** Information of the genomes added to Kraker2 database for the metagenomic analysis.

| Accession | Organims | TaxID |
|---|---|---|
| GCF_001645765.1 | *Candidatus* Mycoplasma haemobos | 432608 |
| GCF_902712995.1 | *Candidatus* Mycoplasma haemohominis | 1494318 |
| GCF_000281235.1 | *Candidatus* Mycoplasma haemolamae str. Purdue | 1212765 |
| GCF_000319365.1 | *Candidatus* Mycoplasma haemominutum 'Birmingham 1' | 1116213 |
| GCF_007858515.1 | *Mycoplasma anserisalpingitidis* | 519450 |
| GCF_000733865.1 | *Mycoplasma buteonis* | 171280 |
| GCF_000012765.1 | *Mycoplasma capricolum subsp. capricolum* ATCC 27343 | 340047 |
| GCF_024918975.1 | *Mycoplasma cottewii* | 51364 |
| GCF_000025845.1 | *Mycoplasma crocodyli* MP145 | 512564 |
| GCF_000687815.1 | *Mycoplasma elephantis* ATCC *51980* | 1408459 |
| GCF_025779955.1 | *Mycoplasma enhydrae* | 2499220 |
| GCF_000327395.2 | *Mycoplasma feriruminatoris* | 1179777 |
| GCF_000238995.1 | *Mycoplasma haemocanis* str. Illinois | 1111676 |
| GCF_000200735.1 | *Mycoplasma haemofelis* str. Langford 1 | 941640 |
| GCF_024722375.1 | *Mycoplasma iguanae* | 292461 |
| GCF_000518305.1 | *Mycoplasma imitans* ATCC 51306 | 1399794 |
| GCF_000253095.1 | *Mycoplasma leachii* 99/014/6 | 866629 |
| GCF_000622205.1 | *Mycoplasma leonicaptivi* ATCC 49890 | 1448135 |
| GCF_004335975.1 | *Mycoplasma marinum* | 1937190 |
| GCF_013008635.1 | *Mycoplasma miroungigenitalium* | 754515 |
| GCF_013008815.1 | *Mycoplasma miroungirhinis* | 754516 |
| GCF_000023685.1 | *Mycoplasma mycoides subsp. capri* str. GM12 | 436113 |
| GCF_006228185.1 | *Mycoplasma nasistruthionis* | 353852 |
| GCF_000508245.1 | *Mycoplasma ovis* str. Michigan | 1415773 |
| GCF_000477415.1 | *Mycoplasma parvum* str. Indiana | 1403316 |
| GCF_012934855.1 | *Mycoplasma phocoenae* | 754517 |
| GCF_012934885.1 | *Mycoplasma phocoeninasale* | 2726117 |
| GCF_017052595.1 | *Mycoplasma procyoni* | 568784 |
| GCF_900476175.1 | *Mycoplasma putrefaciens* | 2123 |
| GCF_000702705.1 | *Mycoplasma simbae* ATCC 49888 | 1408469 |

| | | |
|---|---|---|
| GCF_003855455.1 | *Mycoplasma struthionis* | 538220 |
| GCF_000203215.1 | *Mycoplasma suis* KI3806 | 708248 |
| GCF_016925555.1 | *Mycoplasma tauri* | 547987 |
| GCF_004362335.1 | *Mycoplasma testudineum* | 244584 |
| GCF_000687795.1 | *Mycoplasma testudinis* ATCC 43263 | 1408471 |
| GCF_004335995.1 | *Mycoplasma todarodis* | 1937191 |
| GCF_014068355.1 | *Mycoplasma tullyi* | 1612150 |
| GCF_000277795.1 | *Mycoplasma wenyonii* str. Massachusetts | 1197325 |
| GCF_000875755.1 | *Mycoplasma yeatsii* GM274B | 743967 |
| GCF_025486335.1 | *Mycoplasma zalophi* | 191287 |
| GCF_024742155.1 | *Mycoplasma zalophidermidis* | 398174 |
| GCF_002290085.1 | *Mesoplasma chauliocola* | 216427 |
| GCF_002999455.1 | *Mesoplasma coleopterae* | 324078 |
| GCF_002930145.1 | *Mesoplasma corruscae* | 216874 |
| GCF_002749675.1 | *Mesoplasma entomophilum* | 2149 |
| GCF_000008305.1 | *Mesoplasma florum* L1 | 265311 |
| GCF_000701525.1 | *Mesoplasma grammopterae* ATCC 49580 | 1408447 |
| GCF_002441935.1 | *Mesoplasma lactucae* ATCC 49193 | 81460 |
| GCF_002804105.1 | *Mesoplasma melaleucae* | 81459 |
| GCF_000702725.1 | *Mesoplasma photuris* ATCC 49581 | 1408448 |
| GCF_000518725.1 | *Mesoplasma seiffertii* ATCC 49495 | 1336238 |
| GCF_002843565.1 | *Mesoplasma syrphidae* | 225999 |
| GCF_002804025.1 | *Mesoplasma tabanidae* | 219745 |
| GCF_000483165.1 | *Acholeplasma multilocale* ATCC 49900 | 1278299 |
| GCF_002930155.1 | *Entomoplasma ellychniae* | 2114 |
| GCF_002804205.1 | *Entomoplasma freundtii* | 74700 |
| GCF_000518285.1 | *Williamsoniiplasma lucivorax* ATCC 49196 | 1399797 |
| GCF_002803985.1 | *Williamsoniiplasma luminosum* | 214888 |
| GCF_002804005.1 | *Williamsoniiplasma somnilux* | 215578 |
| GCF_000687735.1 | *Acholeplasma equifetale* ATCC 29724 | 1408415 |
| GCF_017052655.1 | *Acholeplasma equirhinis* | 555393 |
| GCF_000526235.1 | *Acholeplasma granularum* ATCC 19168 | 1278304 |
| GCF_900660755.1 | *Acholeplasma hippikon* | 264636 |
| GCF_003385765.1 | *Acholeplasma laidlawii* | 2148 |
| GCF_025742995.1 | *Acholeplasma manati* | 591373 |
| GCF_900444665.1 | *Acholeplasma oculi* | 35623 |
| GCF_025446935.1 | *Acholeplasma vituli* | 69473 |
| GCF_003363775.1 | *Spiroplasma alleghenense* | 216931 |
| GCF_000500935.1 | *Spiroplasma apis B31* | 1276258 |
| GCF_001029245.1 | *Spiroplasma atrichopogonis* | 1114980 |
| GCF_001281045.1 | *Spiroplasma cantharicola* | 362837 |
| GCF_008086545.1 | *Spiroplasma chinense* | 216932 |
| GCF_000400935.1 | *Spiroplasma chrysopicola* DF-1 | 1276227 |
| GCF_001886855.1 | *Spiroplasma citri* | 2133 |

| | | |
|---|---|---|
| GCF_002795265.1 | *Spiroplasma clarkii* | 2139 |
| GCF_002237575.1 | *Spiroplasma corruscae* | 216934 |
| GCF_000565175.1 | *Spiroplasma culicicola* AES-1 | 1276246 |
| GCF_000439455.1 | *Spiroplasma diminutum* CUAS-1 | 1276221 |
| GCF_018831625.1 | *Spiroplasma* endosymbiont of *'Nebria riversi'* | 2792084 |
| GCF_902809815.1 | *Spiroplasma* endosymbiont of *Danaus chrysippus* | 2691041 |
| GCF_023846195.1 | *Spiroplasma* endosymbiont of *Lariophagus distinguendus* | 2935082 |
| GCF_003987485.1 | *Spiroplasma* endosymbiont of *Megaselia nigra* | 2478537 |
| GCF_918697755.1 | *Spiroplasma* endosymbiont of *Phyllotreta cruciferae* | 2886375 |
| GCF_001029265.1 | *Spiroplasma eriocheiris* | 315358 |
| GCF_002813555.1 | *Spiroplasma floricola* 23-6 | 1336749 |
| GCF_004379335.1 | *Spiroplasma gladiatoris* | 2143 |
| GCF_001715535.1 | *Spiroplasma helicoides* | 216938 |
| GCF_001274875.1 | *Spiroplasma kunkelii* CR2-3x | 273035 |
| GCF_001267155.1 | *Spiroplasma litorale* | 216942 |
| GCF_005222125.1 | *Spiroplasma melliferum* | 2134 |
| GCF_000517365.1 | *Spiroplasma mirum* ATCC 29335 | 838561 |
| GCF_002865545.1 | *Spiroplasma monobiae* MQ-1 | 1336748 |
| GCF_003339775.1 | *Spiroplasma phoeniceum* P40 | 1276259 |
| GCF_021496725.1 | *Spiroplasma platyhelix* PALS-1 | 1276218 |
| GCF_009866525.1 | *Spiroplasma poulsonii* | 2138 |
| GCF_000565215.1 | *Spiroplasma sabaudiense* Ar-1343 | 1276257 |
| GCF_000400955.1 | *Spiroplasma syrphidicola* EA-1 | 1276229 |
| GCF_009730595.1 | *Spiroplasma tabanidicola* | 324079 |
| GCF_000439435.1 | *Spiroplasma taiwanense* CT-1 | 1276220 |
| GCF_001262715.1 | *Spiroplasma turonicum* | 216946 |
